# Stability of *orb* mRNA in *Drosophila* ovaries is regulated by *cis*-regulatory elements in the 3’ UTR and influences germ cell specification

**DOI:** 10.64898/2025.12.08.693029

**Authors:** Mariya Zhukova, Konstantin V. Yakovlev, Justinn Barr, Rudolf A. Gilmutdinov, Anna Tvorogova, Paul Schedl, Yulii V. Shidlovskii

**Affiliations:** Laboratory of Gene Expression Regulation in Development, Institute of Gene Biology, Russian Academy of Sciences, 119334 Moscow, Russia; Laboratory of Cytotechnology, A.V. Zhirmunsky National Scientific Center of Marine Biology, Far Eastern Branch, Russian Academy of Sciences, Vladivostok, Russia; Department of Molecular Biology, Princeton University, Princeton, NJ 08544-1014, USA; Center for Precision Genome Editing and Genetic Technologies for Biomedicine, Institute of Gene Biology, Russian Academy of Sciences, 119334 Moscow, Russia; Department of Biology and General Genetics, Sechenov First Moscow State Medical University (Sechenov University), 119992 Moscow, Russia

**Keywords:** Orb, CPE motifs, mRNA stability, null mutation

## Abstract

Orb is one of two *Drosophila* mRNA-binding CPEB proteins that activate or repress translation by mediating polyadenylation and deadenylation of target mRNAs through interaction with CPE motifs in their 3’UTRs. Orb is one of the key regulators of oocyte specification and polarity. This regulation acts through establishment and maintenance of a 3’UTR-dependent positive autoregulatory loop. In 16-cell cysts, positive autoregulation is necessary to accumulate high levels of *orb* mRNA and protein in one of the two pro-oocytes. A previous study showed that deletion of the *orb* 3’UTR fragment containing CPE sites leads to a failure in proper *orb* mRNA and protein localization; as a result, oocytes are not specified in 16-cell cysts, and egg chambers contain only nurse cells. In the present study, we found that deletion of CPE and CPE-like motifs disrupts proper Orb protein localization and oocyte specification in germaria. CPE motifs are indispensable for *orb* mRNA stability: deletion of CPE and CPE-like sequences reduces *orb* mRNA stability to one third of level in wild type. When the 3’UTR is deleted, the remaining 115-nucleotide sequence, which does not include CPE motifs, is sufficient for nurse cell development. Removal of this sequence leads to a further decrease in *orb* mRNA stability and a drop of Orb protein to undetectable levels. Oogenesis stops in region 2a/2b of the germarium, where the checkpoint is localized, and egg chambers do not form. We propose that CPE motifs are required for *orb* 3’UTR-mediated Orb accumulation which drives oocyte specification, whereas low levels of unlocalized Orb protein expressed in the absence of Orb autoregulation are sufficient for nurse cell specification.

## Introduction

Cytoplasmic polyadenylation element binding (CPEB) proteins control mRNA localization and translation through their binding to the 3’UTRs of target transcripts. CPEB proteins are components of cytoplasmic ribonucleoprotein complexes that regulate poly(A) tail length of mRNAs, a crucial determinant of transcript stability and translational status. CPEB proteins control local translation of mRNAs when localized in different subcellular domains. Their function is essential for meiotic and mitotic progression, cell metabolism and migration, neuronal plasticity and memory formation. CPEB proteins are also required for germ cell development and early embryogenesis in *Drosophila*, *C. elegans*, *Xenopus* and mouse (Charlesworth et al., 2013; Ivshina et al., 2014; Kozlov et al., 2021). CPEB proteins can act as translational activators or repressors depending on their post-translational modifications. It has been shown that the functional state of CPEB proteins depends on these modifications. In vertebrate and *Drosophila* oocytes, the switch from repressive to activating features of CPEB proteins is regulated by phosphorylation (Atkins et al., 2005; Duran-Arqué et al., 2022; Guillén-Boixet et al., 2016; Hodgman et al., 2001; Huang et al., 2002; Mendez et al., 2000b, 2000a).

All CPEB proteins contain two RRM domains responsible for mRNA binding followed by a zinc-finger domain at the C-terminus. Some proteins also contain poly-Q domains (Charlesworth et al., 2013; Ivshina et al., 2014). The interaction of CPEB proteins with target mRNAs is mediated through recognition and binding to cytoplasmic polyadenylation elements (CPEs) within the 3’UTR. CPEs were first discovered in mRNAs of *Xenopus* oocytes as sequences generally consisting of U_3-5_A_1-3_U (Charlesworth et al., 2004). *Xenopus* CPEB1 can weakly interact with non-consensus sequences such as UUUUACU, UUUUAACA, UUUUAAGU and UUUCAU (Charlesworth et al., 2013; Kozlov et al., 2021). The *Drosophila* genome contains two CPEB-related genes, *orb* and *orb2*. Orb and Orb2 proteins have affinities for different CPE sequences. In S2 cells, Orb preferably binds to consensus CPEs consisting of UUUUA_1-3_U, which are similar to the consensus CPEs of vertebrates (Charlesworth et al., 2004; Tay et al., 2000). Orb2 can bind to both A- and G-containing CPEs (Stepien et al., 2016).

Orb regulates oogenesis at multiple stages, including specification of oocytes in 16-cell cysts and establishment of oocyte anterior-posterior and dorsal-ventral polarity (Barr et al., 2019a, 2024; Castagnetti and Ephrussi, 2003; Christerson and McKearin, 1994; Huynh and St Johnston, 2000; Lantz et al., 1994). Orb is also expressed in testes and in the adult brain, where it participates in long-term memory formation (Lantz et al., 1992; Pai et al., 2013). In oocytes, Orb mediates the localization and on-site translation of target mRNAs. Orb regulates the localization and translation of the polarity determinants *oskar* (*osk*), *fs(1)K10* and *gurken* (*grk*) (Castagnetti and Ephrussi, 2003; Chang et al., 1999, 2001; Davidson et al., 2016). Orb also binds *aPKC* mRNA, which encodes a protein of the anterior polarization complex Par, suggesting that its translational activity is regulated in an Orb-dependent manner (Barr et al., 2019a; Benton and St Johnston, 2003).

A notable feature of Orb is its interaction with *orb* mRNA. Orb regulates the localization of its own mRNA in oocytes and subsequently activates *orb* mRNA translation, leading to the on-site accumulation of Orb protein (Tan et al., 2001). In ovaries, this autoregulatory mechanism is activated in oocytes and repressed in nurse cells (Tan et al., 2001; Wong and Schedl, 2011). The autoregulatory loop also drives oocyte specification in the germaria through an *orb* 3’UTR-dependent mechanism. Deletion of the 3’UTR leads to failure in oocyte specification and all germline cells in cysts become nurse cells (Barr et al., 2019b). A recent study showed that in the germarium, the *orb* 3’UTR associates with a special membranous structure called the fusome, which allows firstly to accumulate *orb* mRNA and then Orb protein in the presumptive oocyte (Barr et al., 2024). The fusome is also required to synchronize the cell divisions of cystocytes in the cyst (De Cuevas et al., 1996).

In this work, we have studied the functional significance of CPE motifs as they are potentially essential for Orb autoregulation. Additionally, we have investigated the effect of the last 115 nucleotides of the *orb* 3’UTR and showed that this sequence is necessary for *orb* mRNA stability and translation after disruption of Orb autoregulatory loop. We found that deletion of 12 CPEs and CPE-like motifs disrupts Orb protein localization in cysts and egg chambers. Oocytes fail to specify, and the egg chambers contain only nurse cells. Deletion of either the main part of the 3’UTR or the CPE motifs within the 3’UTR results in decreased mRNA stability, but these mutants differ in the localization of Orb protein in ovaries. The removal of the last 115 nucleotides remaining after the *orb* 3’UTR deletion causes further decrease of *orb* mRNA stability and Orb protein drops to undetectable level. Similar to *orb* null mutants, 16-cell cysts fail to pass the checkpoint in region 2a/2b of the germaria. We propose that in ovaries Orb can regulate germ cell development via two mechanisms. The Orb autoregulatory loop is essential for oocyte specification, and the CPE motifs are indispensable for establishing and maintaining this machinery, while the development of nurse cells requires Orb but does not depend on *orb* autoregulation.

## Results

### Characterization of generated orb 3’UTRs

The *orb* 3’UTR is an essential element of the Orb-dependent positive autoregulatory loop, which underlies oocyte specification and polarity. As previously shown, *orb-RA* is the major isoform expressed in ovaries, consisting of 4,862 nucleotides of the coding region and a 1,206-nucleotide 3’UTR (Barr et al., 2019b; Christerson and McKearin, 1994). Deletion of nucleotides 24 to 1,091 in the 3’UTR resulted in failure of oocyte specification due to mislocalization of *orb* mRNA and Orb protein. This stock, called *orb^Δ3’UTR^*, was generated by replacing the 24-1,091 nucleotide fragment of the *orb-RA* 3’UTR, which contains 12 CPEs and CPE-like sequences, with a *DsRed* expression cassette (referred to hereafter as the *DsRed* gene). The DsRed gene consists of a 3xP3 promoter, the *DsRed* coding sequence, and an SV40 terminator. The *DsRed* gene was flanked by LoxP sites and was subsequently deleted by Cre-LoxP recombination (Barr et al., 2019b). To restore *orb* function, the *orb^resc^*stock was produced by inserting the 24-1,091-nt fragment of the *orb* 3’UTR back into the *orb^Δ3’UTR^* line. To study the significance of CPEs in *orb* mRNA in the context of CPE-dependent Orb autoregulation, the *orb^ΔCPE^*stock was obtained by inserting a 30-1,169-nt fragment of the 3’UTR with deleted consensus and non-consensus CPEs and CPE-like sequences. To investigate the significance of the last 115 nucleotides of the *orb* 3’UTR, two intermediate stocks with the *DsRed* gene inserted into the 3’UTR were analyzed. The *DsRed* insertion deleted this sequence from the 3’UTR because it is localized downstream of the DsRed and is not transcribed as part of the *orb* 3’UTR. The first stock, *orb^Δ3’UTR-DsRed^*, is an intermediate line before *DsRed* excision to produce *orb^Δ3’UTR^*. In *orb^Δ3’UTR-DsRed^*, the terminal 115-nt sequence is completely deleted. The second stock, *orb^ΔCPE-DsRed^*, represents the intermediate stage of 3’UTR replacement for generating *orb^ΔCPE^*. In this case, the 3’UTR includes 78 nucleotides from the terminal 115-nt sequence, but without PAS.

When *orb^Δ3’UTR^* was generated by CRISPR/Cas9, an additional stop codon was added in the translated region of the last exon during the 3’UTR deletion (Figure 1A). The additional stop codon resulted in the loss of a non-conserved 75-amino acid sequence from the C-terminus and a decrease in the protein’s molecular weight by 8.8 kDa (Figure 1B). Hence, the sequence downstream of the additional stop codon became part of the 3’UTRs in all generated stocks. The shortened Orb remains functional, as a line with a single point mutation *orb^T840stop^* showed only a slight decrease in fecundity (approximately 10%) (Figure S1). *orb^resc^* exhibited a mild decrease in fecundity (∼33%). Females with other 3’UTRs were sterile and did not lay eggs. Visual observations of dissected ovaries showed that the generated 3’UTRs had a negative effect on ovarian size (Figure 1C). Late-stage oocytes and eggs were present in WT and in the smaller *orb^resc^* ovaries. However, in *orb^Δ3’UTR^*, *orb^ΔCPE^* and *orb^ΔCPE-DsRed^*, ovaries did not contain egg chambers at late stages of oogenesis and included only small and mid-size follicles. The most detrimental effect was observed in *orb^Δ3’UTR-DsRed^*. These females had small ovaries with no clear presence of follicles.

**Fig 1.**
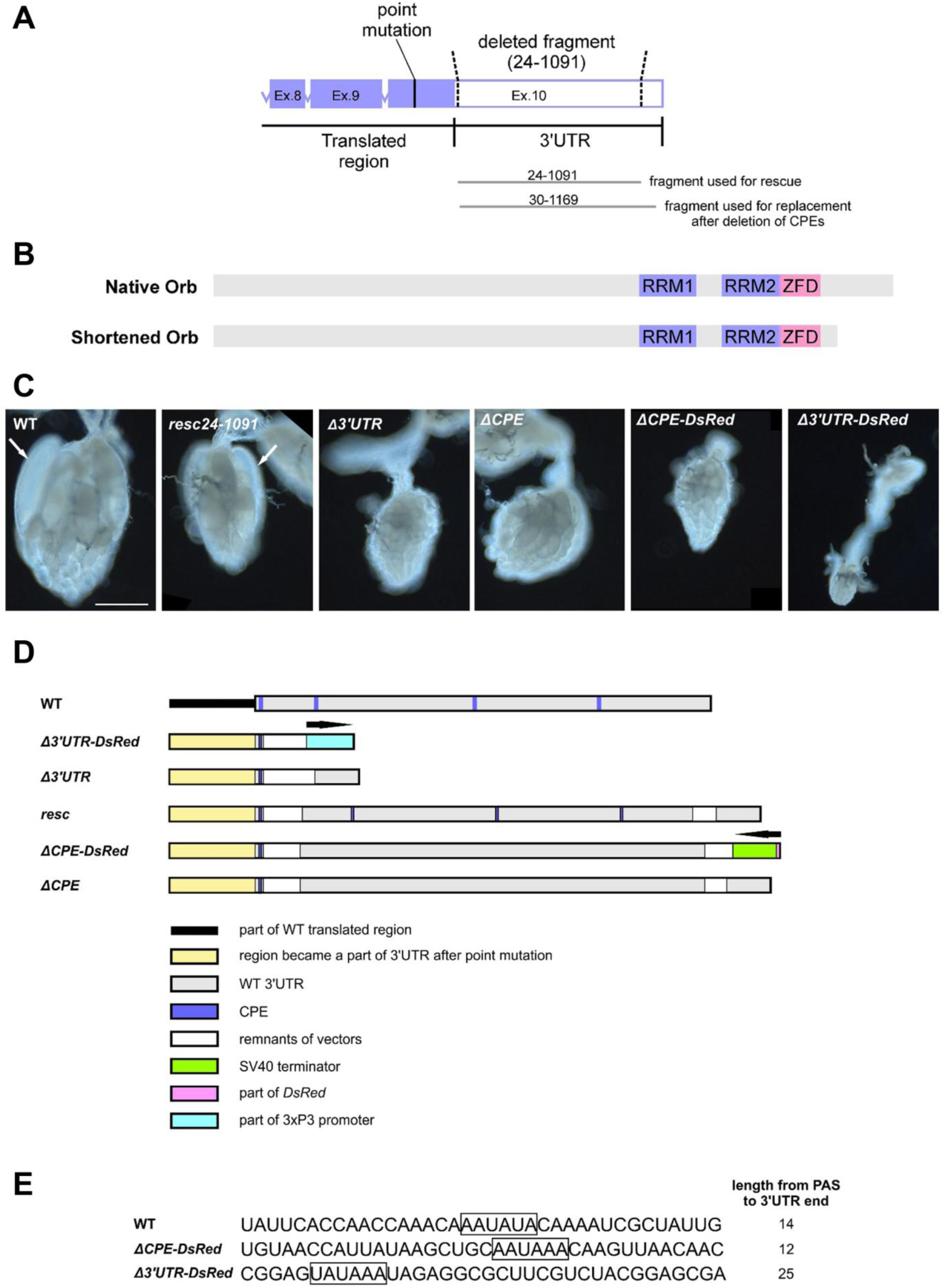
Characterisitic of generated flies in the study. (**A**) Schematic representation of part of the *orb-RA* genomic organization and a strategy of deletion, rescue and replacement of the 3’UTR. Point mutation localizes in the end of the translated region leading to Orb protein shortening. Ex. – exon. (**B**) Scheme of encoded Orb proteins. Comparison of structures of native and shortened Orb. Q-rich domains localized in N-terminus part are not shown. (**C**) *orb* 3’UTR replacements negatively effect on ovarian size. Whole mount images of three-day ovaries were obtained by using dark-field microscopy at the same magnification. Arrows point to eggs. Bar is 0.25 mm. (**D**) A scheme of ovarian *orb* 3’UTRs after genetic manipulations confirmed by 3’-RACE. Schematic sequence organization of the *orb* 3’UTRs after deletion and replacement procedures and in intermediate lines with *DsRed* marker (**E**) PAS hexamer sequences located near 3’UTR ends. WT corresponds to four lines with similar 3’ end: WT, *orb^resc^*, *orb^Δ3’UTR^* and *orb^ΔCPE^*.

A valid analysis of the effect of the obtained 3’UTRs on oogenesis requires precise information about their expressed sequences. To determine whether the transcription termination site in the *orb* 3’ UTR was altered in the new mutants, we performed PCR and 3’-Step-Out RACE. Exact 3’UTR sequences were determined by direct sequencing of PCR fragments (Figure 1E, the sequences of the 3’UTRs are provided in Supplementary File S1). Nuclear cleavage and polyadenylation of pre-mRNA require at least two signals: the core polyadenylation signal (PAS) and the downstream sequence element (DSE). The PAS is located 10-30 nt upstream of the cleavage site (CS), while the DSE is located within the first 30 nt downstream of the CS (Sanfilippo et al., 2017; Tian and Graber, 2012). We identified *Drosophila*-specific PAS sequences located near the 3’ ends in all 3’UTRs (Figure 1E), but did not find evident DSEs in the downstream genomic sequences, as these motifs are less conserved (Retelska et al., 2006; Sanfilippo et al., 2017; Smibert et al., 2012). As expected, the ovarian 3’UTR of *orb* mRNA in *orb^Δ3’UTR^*, *orb^ΔCPE^* and *orb^resc^* had the same 115-nt 3’end as in WT, which remained after the deletion of the 3’UTR 24-1,091 fragment and retained an intact PAS (Figure 1E). In the genomes of *orb^Δ3’UTR-DsRed^* and *orb^ΔCPE-DsRed^*, the 115-nt sequence of the WT 3’UTR was located downstream of the *DsRed* gene. The presence of *DsRed* in both the direct and reverse orientations in the 3’UTR led to the formation of new 3’ ends that included parts of the *DsRed* gene (Figure 1D). The new pre-mRNA included different PAS motifs (Retelska et al., 2006) and remnants of the inserted vector upstream of them. In *orb^Δ3’UTR-DsRed^*, the original AAUAUA hexamer was changed to UAUAAA, while in *orb^ΔCPE-DsRed^*ovaries, the original hexamer was changed to AAUAAA (Figure 1E).

### Orb protein and mRNA expression in ovaries with different 3’UTRs

It was previously shown that deletion of the *orb* 3’UTR disrupts *orb* mRNA and protein localization and high-level expression of Orb protein that leads to the failure of oocyte specification and hence female sterility (Barr et al., 2019b). Our observations showed that replacement of the last 115 nt of the *orb* 3’UTR with a non-functional sequence in the *orb^Δ3’UTR-DsRed^* and *orb^ΔCPE-DsRed^* lines also led to female sterility. In both cases, females had small ovaries with no visible eggs. To elucidate how 3’UTR mutations influence oogenesis, we first examined Orb protein expression. Due to the generation of a new stop codon, ovaries with mutated 3’UTRs expressed truncated Orb protein, which was smaller than in WT (Figure 2A). The shortened Orb appeared as two bands, similar to the WT. The upper and lower bands corresponded to the slow-migrating hyperphosphorylated form and the fast-migrating hypophosphorylated form of the protein, respectively (Wong et al., 2011). To examine Orb mobility in polyacrylamide gel, we analyzed *orb-XN* 3’UTR ovaries, which expressed the shortened Orb. After treatment with λ protein phosphatase, both native and shortened Orb were detected as a single fast-migrating band (Fig. 2B). This confirms our suggestion that deletion of the C-end fragment by generation of the new stop codon does not affect Orb phosphorylation, as none of the predicted phosphorylation sites are located in the deleted protein sequence (Wong et al., 2011). In *orb^Δ3’UTR-DsRed^* ovaries, no Orb signal was detected even after overexposure (Figure S2).

**Fig 2.**
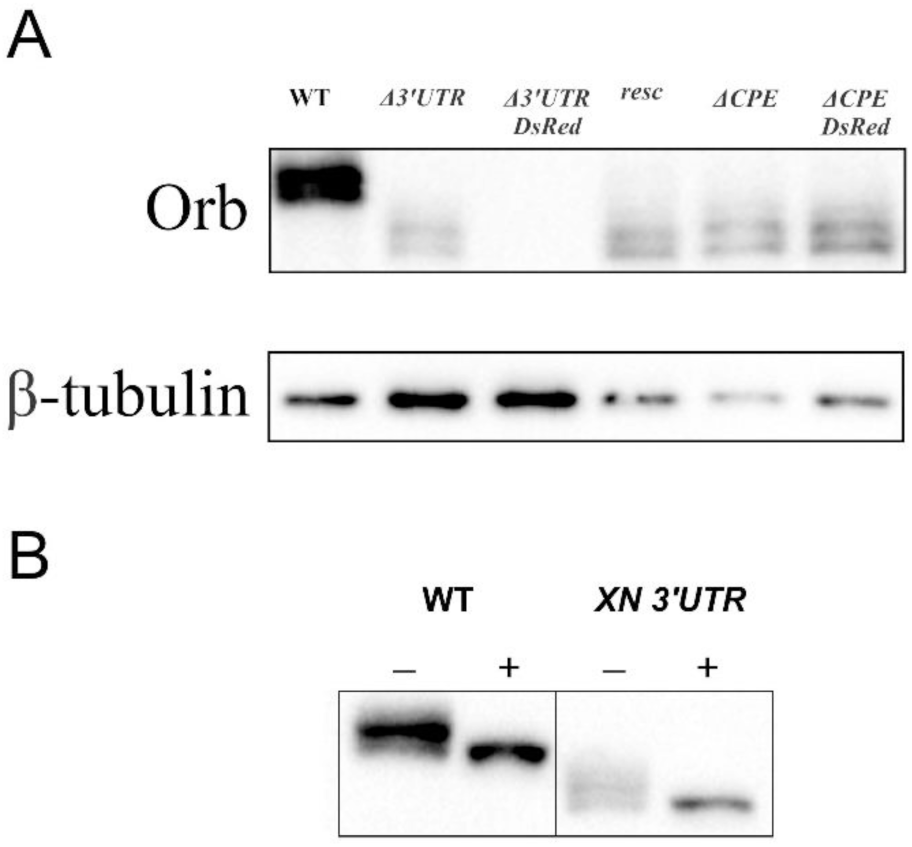
Western blot analysis of Orb in 3-5-day ovaries. (**A**) Orb is detected as two bands. In ovaries with mutated *orb* 3’UTRs Orb migrates faster than WT. β-tubulin was used as loading control. (**B**) Ovarian samples treated by λ protein phosphatase. After treatment (+), native Orb in WT and shortened Orb in *orb-XN* 3’UTR are detected as single fast migrated band.

Our results showed that the generated *orb* 3’UTRs negatively affect the amount of Orb protein in ovarian germ cells. Next, we measured the relative amount of *orb* mRNA in the ovaries. According to our RT-qPCR results, the mRNA level in fertile females after 3’UTR rescue in *orb^resc^* was half of the level in WT (Figure 3A). At the same time, the levels in WT and *orb^ΔCPE^* were the same and the level of *orb^Δ3’UTR^*was slightly higher than in *orb^resc^*. In *orb^Δ3’UTR-DsRed^*, the mRNA level was extremely low (0.036 of WT level), which explains the undetectable level of Orb protein due to the very low amount of translated *orb* mRNA in these females. These results did not correlate with the fecundity of the corresponding flies, as sterile *orb^ΔCPE^*, *orb^Δ3’UTR^* and *orb^ΔCPE-DsRed^* flies had higher levels of *orb* mRNA than the fertile *orb^resc^*.

**Fig 3.**
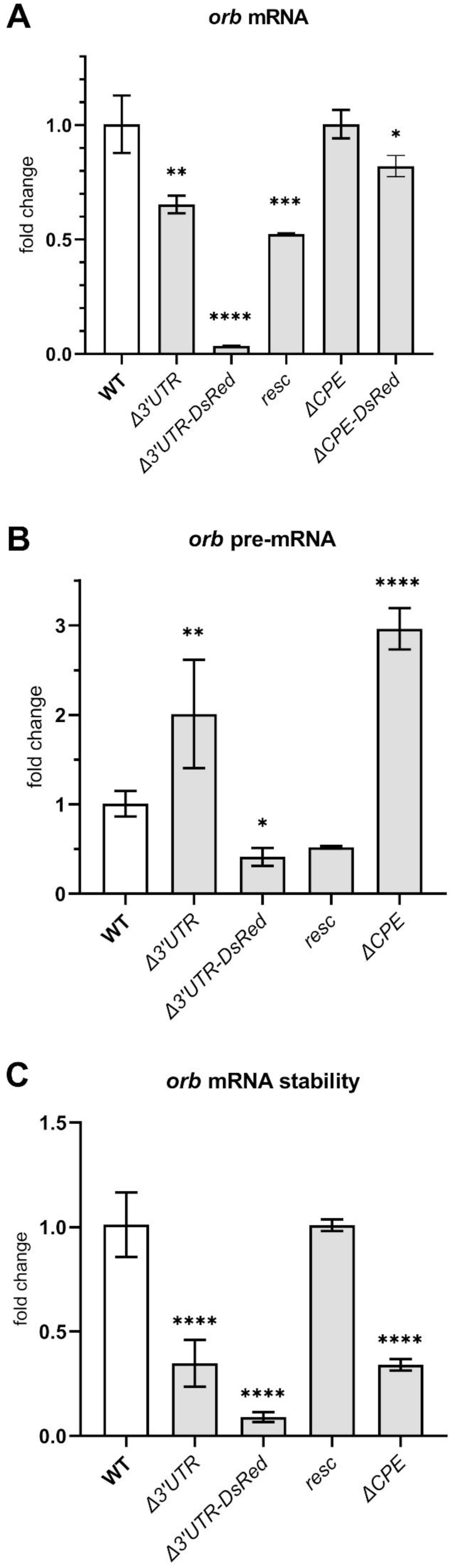
Expression of *orb* in ovaries estimated by RT-qPCR. (**A**) Expression of *orb* mRNA (n=2). (**B**) Evaluation of the *orb* transcriptional levels by measurement of *orb* pre-mRNA (n=3). (**C**) Levels of the *orb* mRNA stability calculated by ratio of relative level of mRNA to relative level of pre-mRNA. Values are shown as mean ± SD. Statistical significance estimated by comparison test with WT is shown. * p < 0.05, ** p < 0.01, *** p < 0.001, **** p < 0.0001.

We hypothesized that the significant fluctuations in *orb* mRNA levels between flies could be related to post-transcriptional regulation of mRNA. Particularly, the diversity in mRNA levels might be associated with differences in mRNA stability, which depends on the 3’UTR sequence. To test this hypothesis, we measured the level of *orb* nascent pre-mRNA and used these values to calculate the mRNA stability coefficient. As shown in Figure 3B, the *orb* pre-mRNA level was elevated in both *orb^Δ3’UTR^*and *orb^ΔCPE^* (with two-fold and three-fold increase, respectively). In contrast, the *orb* pre-mRNA level decreased in *orb^Δ3’UTR-DsRed^*(0.41). A decreased level was also found in *orb^resc^* (0.52), although this value did not differ significantly from WT.

Next, we estimated the relative mRNA stability of *orb* by calculating the ratio of mRNA to pre-mRNA (Katsioudi et al., 2023; Kortenoeven et al., 2012). A lower mRNA/pre-mRNA ratio compared to WT indicates a decrease in mRNA stability, and vice versa. We examined mRNA/pre-mRNA ratios by dividing the mean mRNA level by the mean of pre-mRNA levels (Figure 3C). The *orb* mRNA stability in WT was set to 1, as both pre-mRNA and mRNA levels were designated as 1. Deletion of either the 3’UTR in *orb^Δ3’UTR^* or the CPE motifs in *orb^ΔCPE^*led to a three-fold decrease in *orb* mRNA stability. The strongest negative effect was observed in *orb^Δ3’UTR-DsRed^* ovaries, where the stability level dropped by more than ten-fold. In *orb^resc^*ovaries, the level of mRNA stability was fully restored to the WT level. We found that *orb* mRNA stability depends on the sequence of its 3’UTR. Specifically, either deletion of the 3’UTR or CPE sites resulted in nearly the same *orb* mRNA/pre-mRNA ratios. It means that CPE sites are involved in control of *orb* mRNA turnover by regulating mRNA stability. The last 115 nucleotides in the 3’UTR are also essential for *orb* mRNA stability, as the substitution of this sequence with the inserted vector (Figure 3A) led to a further decrease in mRNA stability from 0.33 in *orb^Δ3’UTR^* to 0.09 in *orb^Δ3’UTR-DsRed^*. Our data from the *orb^915Δ3’UTR^*and *orb^915Δ3’UTR-DsRed^* lines are consistent with the results from *orb^Δ3’UTR^* and *orb^Δ3’UTR-DsRed^* (Figure S3, S4).We cannot exactly explain the reasons for the differences in the pre-mRNA levels between strains. It is possible that there are verying numbers of *orb*-expressing cells in ovaries with different 3’UTR mutations or that the mutations in the 3’ UTR directly or indirectly affect *orb* transcription to different degrees.

### Orb localization and oocyte specification in germaria

Oocyte specification depends on the proper localization and accumulation of *orb* mRNA and protein, which in turn depend on the *orb* 3’UTR. Our analysis of *orb* mRNA distribution in the germaria using single-molecule FISH showed that in mutant flies (except *orb^resc^* and *orb^T840stop^*) the mRNA levels were indistinguishably low and close to the background signal (not shown). Therefore, to define the effect of 3’UTR sequences on Orb protein localization and germ cell fate we analyzed ovaries after Orb immunostaining. In early oogenesis, Orb is distributed equally in germ cell cysts and begins to accumulate in two pro-oocytes in region 2a and then in one pro-oocyte, which assumes an oocyte identity in region 2b. Deletion of the 3’UTR leads to failure of oocyte specification and all germ cells become nurse cells with diffuse Orb localization (Barr et al., 2019b). In our experiments, Orb signal was specific in WT pro-oocytes in the germaria (Figure 4A, C, D). The deletion of CPE sites in *orb^ΔCPE^* caused a disruption in proper Orb localization. In region 2a, most cystocytes exhibited an even distribution of Orb, although in some cases two cystocytes with a more intense signal could be distinguished (Figure 4E). In region 2b Orb protein could be distributed evenly, accumulated vaguely between two pro-oocytes (Figure 4 E, F) and in rare cases, the signal was brighter in one pro-oocyte (Figure 4 B, G). In region 3 (or stage 1 of oogenesis) the intensity of the Orb signal was slightly brighter in 1-3 cells, some of which had oocyte-like nuclei. Deletion of the 3’UTR in *orb^Δ3’UTR^*resulted in a faint diffuse Orb signal in the germaria (Figure 4H). In *orb^Δ3’UTR-DsRed^*, Orb was undetectable after immunostaining. Moreover, egg chambers were indiscernible by DAPI staining, suggesting that germ cells had stopped developing in the germaria at the cyst stage (Figure 4I).

**Fig 4.**
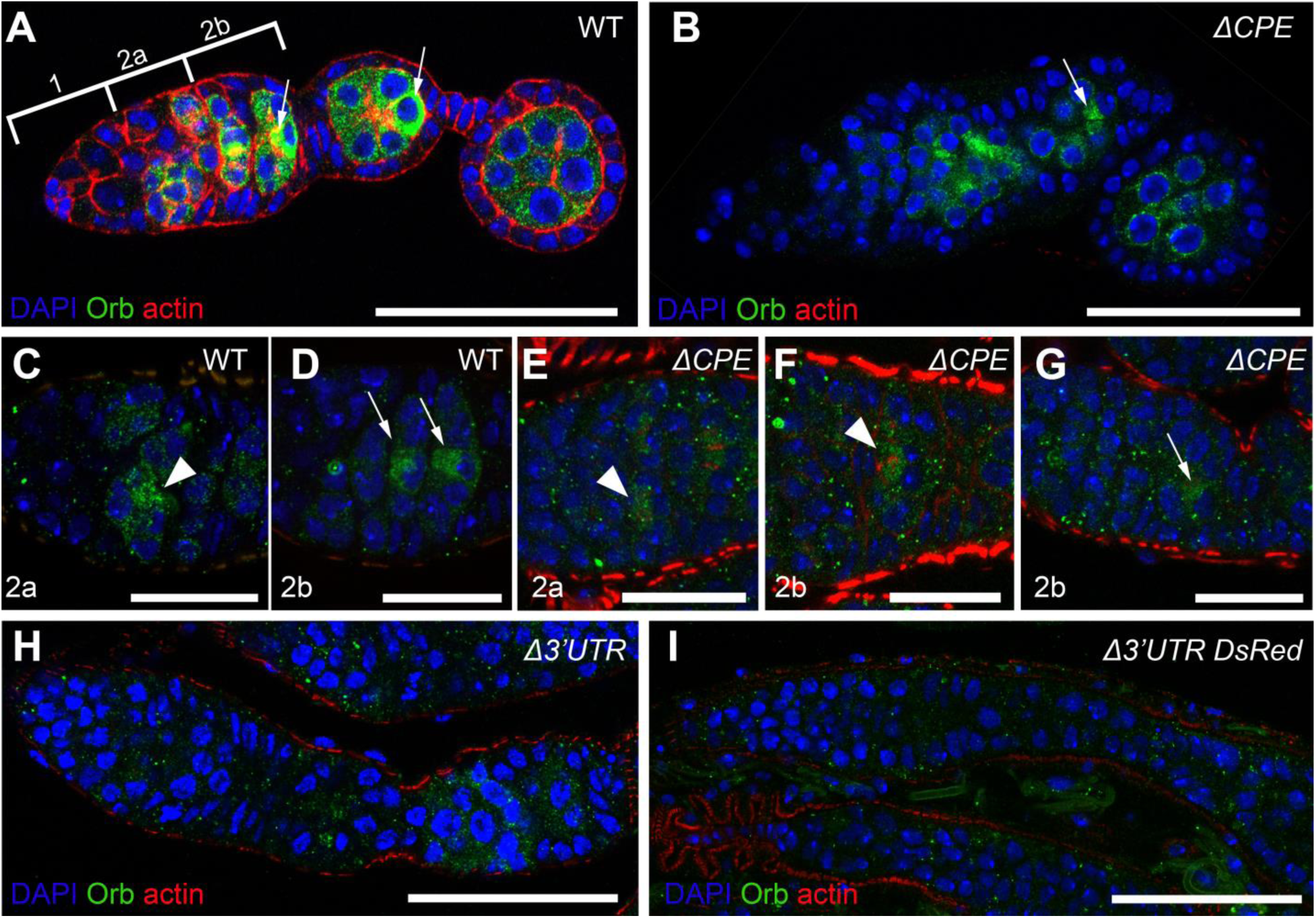
Orb localization in germaria and stage 1 egg chambers. Orb staining is shown in green. Counterstaining was done with DAPI (blue) and Alexa Fluor 633 phalloidin (red). In WT (**A**), oocytes are recognized by concentrated Orb staining. Arrows indicate accumulated Orb in the oocyte. Regions 1, 2a and 2b of the germarium are indicated by brackets. In *orb^ΔCPE^* (**B**), Orb is distributed without clear oocyte-specific localization. In region 3 of the germarium, arrow points to nucleus morphologically similar to those of oocyte. (**C-G**) – accumulation of Orb in germaria at stages 2a and 2b in WT (**C, D**) and *orb^ΔCPE^*(**E-G**). Arrowhead points to two cystoblasts with similarly accumulated Orb protein in the cytolplasm. In *orb^Δ3’UTR^* (**H**), germarium and stage 2 have low Orb amount. Orb is not detected in *orb^Δ3’UTR-DsRed^*germaria (**I**). Bars are 50 µm.

### Cyst development in germaria of mutant flies

To analyze morphological changes in cyst development at early stages of oogenesis, we observed cysts in the germaria after alpha-spectrin immunostaining. Alpha-spectrin is a membrane skeletal protein that is a constituent of the fusome in the germaria and forms the submembrane cytoskeleton of epithelial cells, including follicular cells in the ovaries of *Drosophila* (Lee et al., 1997; Lin et al., 1994). There were no morphological changes in the cysts of the germaria in *orb^Δ3’UTR^*, and the egg chambers appeared normal despite the absence of specified oocytes (Fig. 5B). In *orb^ΔCPE^*, most germaria did not differ from WT, but in some ovaries appeared germaria with disruptions in region 2b. In these germaria, cystocytes of two cysts merged, forming egg chambers with more than 16 cells (20 cells at Fig. 5D). In other germaria, no encapsulated cysts were observed within the layer of follicular cells, and region 2b was entirely composed of follicular cells (Fig. 5E). There were no germaria with normal morphology in *orb^Δ3’UTR-DsRed^*, and all of them had morphological defects. Most germaria lacked encapsulated cysts in region 2b and were extended by a stack of follicular cells without egg chambers (Fig 5F). If an egg chamber was formed, it often consisted of more or fewer than 16 cyst cells. Notably, WT germaria usually contain 2-3 germline stem cells with a round spectrosome, which is a precursor of the fusome, 1-2 cystoblasts, and 2-cell cysts with a fusome either in one cell or spread between two cells. This amounts to five to six cells in total. In *orb^Δ3’UTR-DsRed^*, the number of cells with round-shaped fusome or fusome spread between two cells was often increased to 10-16 cells in region 1 (Fig 5 G, H).

**Fig. 5.**
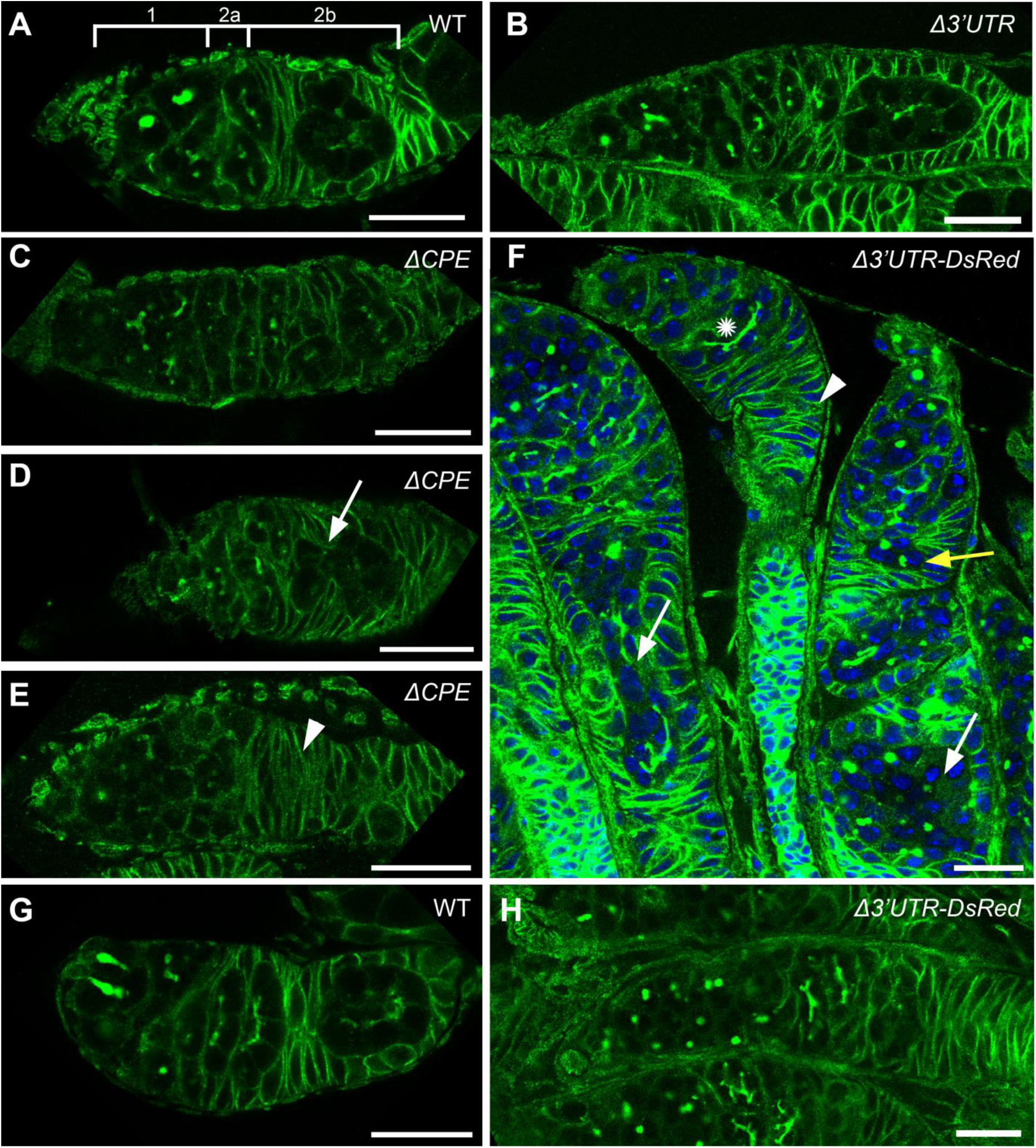
Egg chamber formation in germaria of ovaries. A-E – immunostaining of alpha-spectrin is shown (green), F – immunostaining of alpha-spectrin (green) combined with DAPI staining (blue). A-C – normal morphology of germarium in WT, *orb^Δ3’UTR^* and *orb^ΔCPE^*. Regions 1, 2a and 2b of the germarium indicated by a bracket. D-F – disruptions of egg chamber formation in flies *orb^ΔCPE^* and *orb^Δ3’UTR-DsRed^*. Egg chambers with more than 16 cells indicated by white arrows (D, F), a stack of follicular cells without developing egg inside marked by arrowhead, an egg chamber consisting of five cells pointed by yellow arrow, asterisk marks a normally developed cyst with 16 cells. G – two dividing spectrosomes in the region 1 of the germarium in WT. H – more than 10 round-shaped fusomes in cystocytes in the region 1 of the germarium of *orb^Δ3’UTR-DsRed^*. Bars are 20 µm.

### Orb localization in ovarioles of flies with different orb 3’UTRs

Further examination of Orb localization in later developmental stages also revealed distinct distribution patterns among the lines. In early and mid-oogenesis, Orb predominantly localized in oocytes of each egg chamber of WT ovarioles (Figure 6A). Similar pattern was observed in ovaries of *orb^T840stop^* and *orb^resc^*(Figure S5, S6). Egg chambers of *orb^ΔCPE^* usually contained nurse cells with Orb accumulated in perinuclear area and in clamp-like structures, which resemble perinuclear P bodies (Figure 6B). Some chambers contained 1-3 cells resembling oocytes by condensed nuclei, Orb staining level could be similar to nurse cells or to a small degree higher. About 63% of ovarioles *orb^ΔCPE^* had at least one egg chamber with condensed nuclei, similar to oocyte. Egg chambers in *orb^Δ3’UTR^* did not have oocytes and consisted of nurse cells with low-intensity diffuse Orb staining (Figure 6C), though single condensed nuclei in egg chambers were infrequently observed (Barr et al., 2019b). Besides Orb localization, another difference between WT and *orb^ΔCPE^*/*orb^Δ3’UTR^*egg chambers was observed in nurse cells. Nuclei of WT nurse cells had more uniform staining of DNA by DAPI than in *orb^ΔCPE^*/*orb^Δ3’UTR^*, whose nuclei were more heterogenous. It may indicate that along with oocyte specification the *orb* mRNA 3’UTR is also necessary for functioning of nurse cells, particularly at transcriptional level. The largest egg chamber size in *orb^ΔCPE^*and *orb^Δ3’UTR^* corresponded to mid-oogenesis in WT. It suggests that egg chambers of *orb^Δ3’UTR^* and *orb^ΔCPE^*without oocytes or even with defective oocytes could not further develop and die. In mid-oogenesis, the second checkpoint at stage 8 can activate apoptosis of germ cells in response to poor diet, ecdysone stimuli, abnormal egg chamber development and chemical treatment. Apoptotic cells are easily visible by highly condensed nuclei within egg chambers (McCall, 2004; Nezis et al., 2000; Peterson et al., 2003). Indeed, we detected apoptotic cells in *orb^Δ3’UTR^* egg chambers in distal part of ovarioles after DAPI staining (Figure 6C) that confirm our suggestion. The strongest negative effect was found in *orb^Δ3’UTR-DsRed^* (Figure 6D). As in Western blot analysis, in which we did not detect Orb protein, immunostaining was also close to background level. Also, *orb^Δ3’UTR-DsRed^* ovaries did not contain egg chambers. The ovaries of *orb^915Δ3’UTR^* and *orb^915Δ3’UTR-DsRed^*lines had similar morphology to that described above for *orb^Δ3’UTR^* and *orb^Δ3’UTR-DsRed^*, respectively (Figure S6).

**Fig 6.**
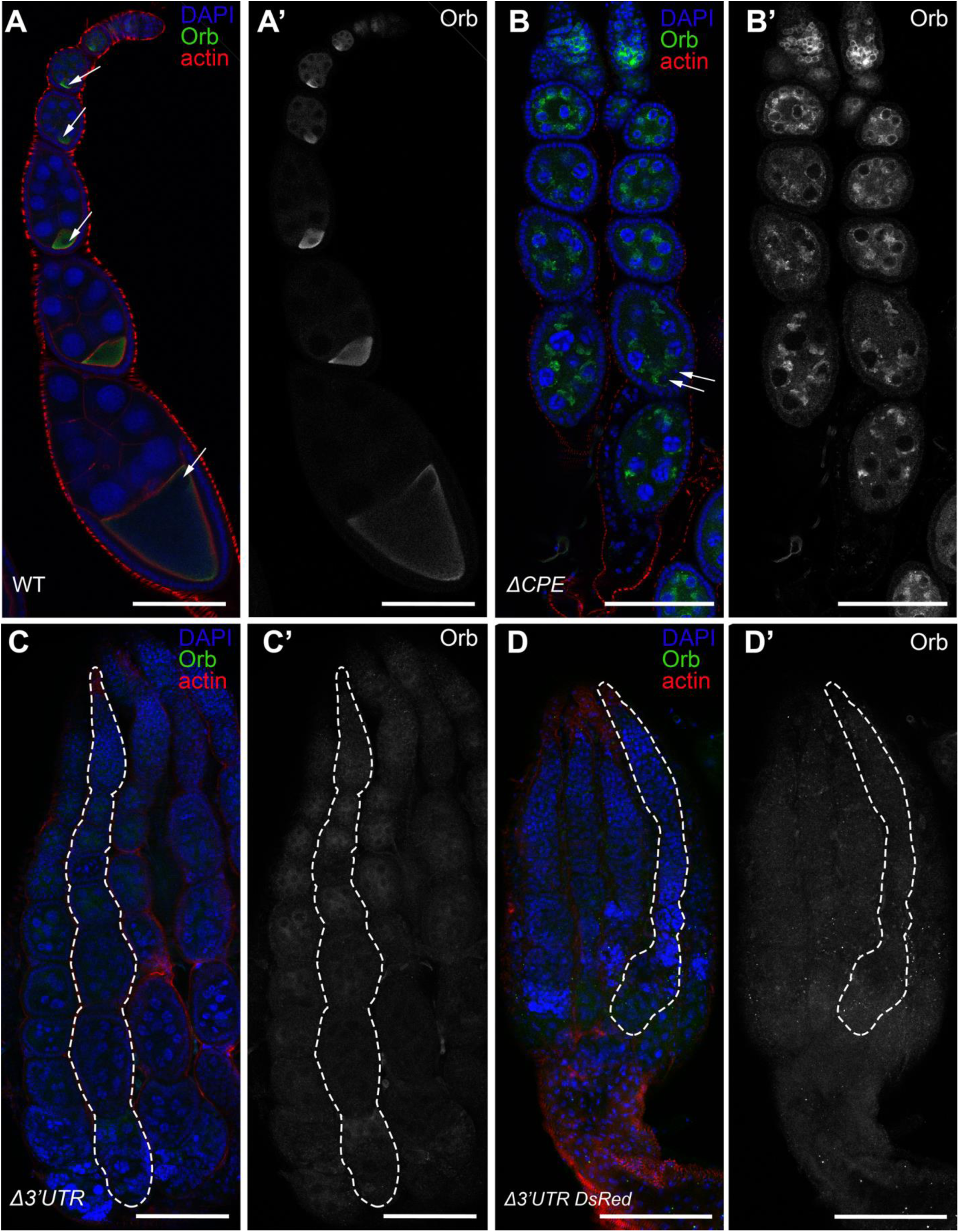
Orb localization in ovaries of flies with different *orb* 3’UTRs. (**A-D**) Ovaries after Orb (green) immunostaining and counterstaining with DAPI (blue) and Alexa Fluor 633 phalloidin (red). Separate images of channel with Orb staining are presented on the right (**A’-D’**). In WT (**A**, **A’**), Orb localizes in oocyte-specific manner in each egg chamber. Arrows indicate nuclei of the oocytes. In *orb^ΔCPE^* (**B**, **B’**), oocytes are not visible by Orb staining, though in one chamber pair of nuclei resemble those in oocytes (double arrow). Major Orb signal is seen in perinuclear area of the nurse cells. In *orb^Δ3’UTR^* (**C**, **C’**), oocytes are absent and a weak diffuse Orb signal localizes in cytoplasm of nurse cells. In *orb^Δ3’UTR-DsRed^* (**D**, **D’**), egg chambers are not visible. Ovarioles are thin and without Orb signal. Bars are 100 µm.

### Expression and localization of germ cell marker vasa

The deletions of ether 3’UTR or CPE sites gave rise to abnormalities in Orb functioning and consequent disturbance of the oocyte specification. More pronounced negative effect was in *orb^Δ3’UTR-DsRed^*, when Orb was not expressed at the protein level and germ cells could not be recognized by Orb localization. Alternatively, germ cells can be visualized by expression of other germ cell specific proteins. To check the presence of germ cells in *orb^Δ3’UTR-DsRed^* ovaries and confirm germ cell identity of Orb-positive cells in other lines, we performed Vasa immunostaining. Vasa is known to be germ cell specific protein expressed in germline, from germline stem cells to nurse cells and oocytes of *Drosophila* (Findley et al., 2003; Hay et al., 1988; Lasko and Ashburner, 1988, 1990). In WT, Vasa was found in cysts in germaria (Styhler et al., 1998). In later developmental stages, Vasa was detected within egg chambers in nurse cells and oocytes until late oogenesis (Figure 7A). Since this time, Vasa was visible in nurse cells (Figure 7A) and in posterior region of the oocyte cytoplasm (not shown). The level of Vasa was lower in both *orb^Δ3’UTR^* and *orb^ΔCPE^*. Vasa localized in cells in germaria and in egg chambers, but without perinuclear concentration (Figure 7B, C). In *orb^Δ3’UTR-DsRed^*, Vasa localized in cyst cells. Cells in germaria resembled typical structure, while egg chambers had different shape, round or elongated, and number of cells in egg chambers varied. Some cyst cells detached from cysts as single cells or as few-cell aggregates (Figure 7D).

**Fig 7.**
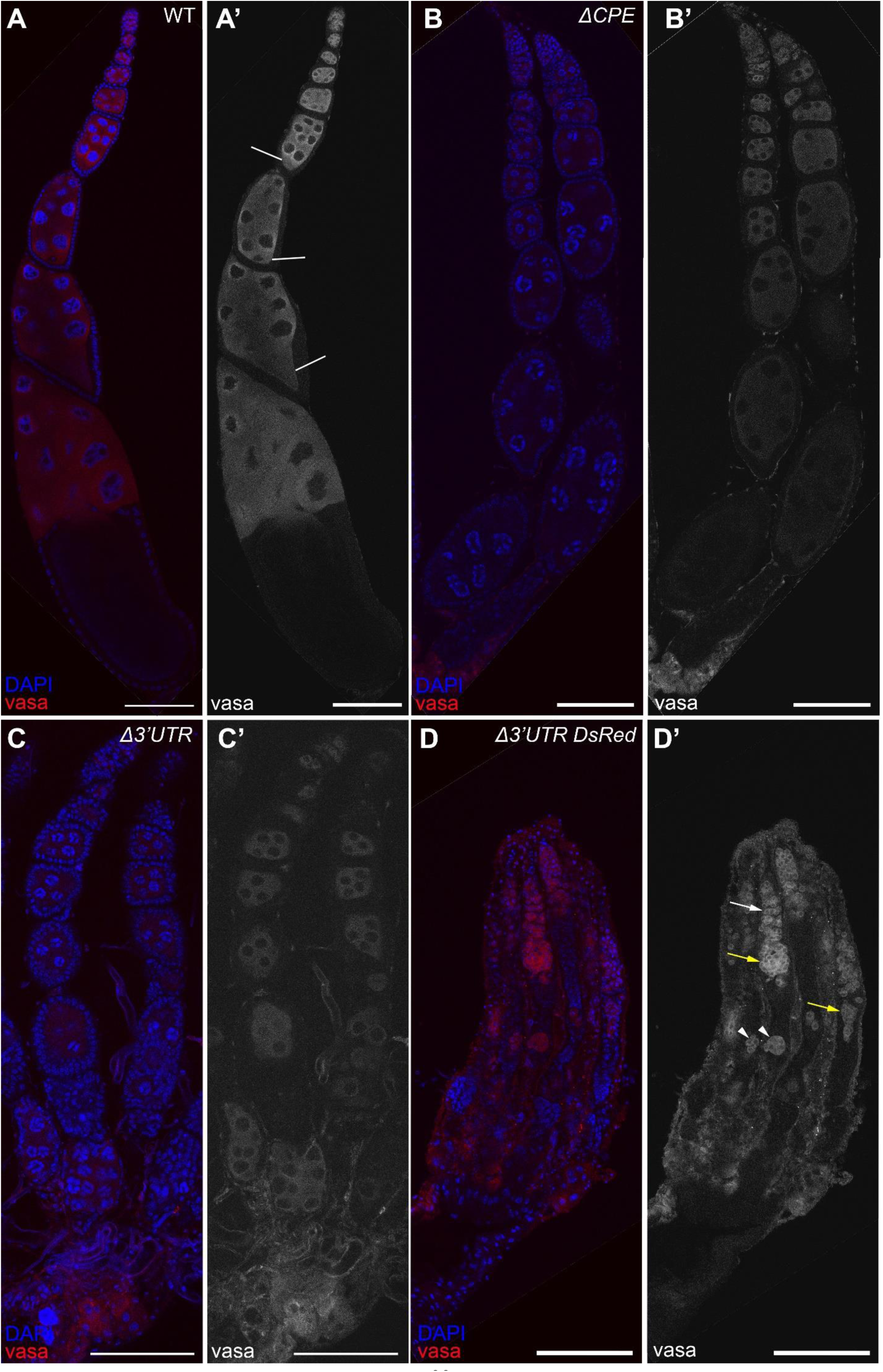
Germ cell visualization by vasa immunostaining. (**A-D**) Vasa protein staining (red) is combined with counterstaining with DAPI (blue). Vasa channel is presented separately on the right (**A’-D’**). (**A**) WT ovaries. Oocytes are marked by lines. In *orb^ΔCPE^* (**B**) and *orb^Δ3’UTR^*(**С**) ovaries, Vasa localizes in germ cell cysts in germaria and differentiated nurse cells in egg chambers, while the level of the Vasa signal is lower than in WT. In *orb^Δ3’UTR-DsRed^* (**D**), Vasa is detected predominantly in proximal part of ovarioles. Unlike in other orb 3’UTRs, germ cells do not develop beyond germ cell cyst and egg chambers do not appear. Some cysts in germaria look normal (arrow), while many egg chambers have abnormal structure (yellow arrows). Some cyst cells are detected in groups or as single cells in more distal parts of the ovarioles (arrowheads). Bars are 100 µm.

As we previously showed, Vasa protein localized in ovarian germ cells in the all *orb* 3’UTR variants. To confirm *vasa* expression on mRNA level, we performed RT-qPCR (Figure S7). *orb^ΔCPE-DsRed^* and *orb^resc^* had a little effect on *vasa* level, which was slightly lower than in WT without statistical significance. Among the 3’UTR mutants, both *orb^Δ3’UTR^* and *orb^ΔCPE^* showed the most considerable decrease of *vasa* level (0.55 and 0.62, respectively). Inversely, in *orb^Δ3’UTR-DsRed^ vasa* level was slightly higher than in WT.

## Discussion

Localized Orb protein is required for oocyte specification and the establishment of oocyte polarity. Orb localization in presumptive and specified oocytes is governed by *orb* mRNA accumulation and subsequent on-site translation. Early studies showed that an 815-nt fragment of the *orb* 3’UTR defined by the HindIII and NdeI restriction sites localized *lacZ* mRNA spatially and temporally resembling the *orb* mRNA pattern (Lantz and Schedl, 1994). Deletion of a major part of the *orb* 3’UTR by CRISPR/Cas9 directly demonstrated the requirement of the 3’UTR-dependent autoregulatory loop for oocyte specification (Barr et al., 2019b). Replacement with partially functional fragment XN (425-815 nt) of the 3’UTR led to partial recovery of oocyte specification, but oocyte polarization remained impaired at later stages (Barr et al., 2019b; Lantz and Schedl, 1994). It was suggested in these studies that proper Orb function throughout oogenesis is maintained by an autoregulatory mechanism involving multiple CPE motifs in the *orb* 3’UTR.

Each CPEB protein has its own repertoire of CPE sites that they bind. CLIP (crosslinking immunoprecipitation) sequencing assay showed that the preferred target sequences of transgenic Orb overexpressing in *Drosophila* S2 cells are linear forms of consensus A-containing CPEs in 3’UTRs. Orb can also bind less frequently to G- and AC-containing CPEs (Stepien et al., 2016). To examine the significance of CPEs for Orb function, we deleted CPEs from the 3’UTR by replacement. Considering the variability of CPE sites that Orb binds to, we removed potential A-, G- and AC-containing CPEs from the *orb* 3’UTR. Our results provide evidence that CPE sites are necessary for stability of *orb* mRNA, as it decreased significantly in *orb^ΔCPE^* flies.

Our detailed analysis of oocyte specification in *orb^ΔCPE^*showed that CPE-like motifs are essential not only for mRNA stability, but also for the transport and activation of the *orb* autoregulatory loop. In the germaria of WT flies, Orb protein is weakly enriched in two pro-oocytes in 16-cell cysts in region 2a. A similar Orb localization was observed in *orb^ΔCPE^* at this stage. The fusome-associated transport activates during cyst transition through region 2a and directs BicD:Egl:*orb*mRNA complexes to accumulate *orb* mRNA in two pro-oocytes The *orb* mRNA transport and autoregulatory regulation of *orb* expression lead to the concentration of Orb protein in one pro-oocyte or in two cells with a clear “winner” in region 2b (Barr et al., 2024). In *orb^ΔCPE^*, Orb accumulation was significantly weaker and it could be observed in all cells of the cyst, in two or in one pro-oocytes, suggesting that transport of *orb* mRNA was not properly activated. This disruption in transport appears to be related to the overlap of two CPE-like motifs which were removed with the K10 transport/localization sequence, that is recognized by Egl transport protein (Bullock et al., 2010). The decreased Orb signal in pro-oocytes implies that the autoregulatory loop for translational activation depends on Orb binding to the removed CPE sites.

Differences in Orb protein staining levels suggest that other sites retained in *orb^ΔCPE^* but deleted in *orb^Δ3’UTR^*are necessary for more efficient Orb translation. Also, *orb^ΔCPE^*has motifs in the 3’UTR required for Orb intracellular localization in nurse cells, as Orb localization in egg chambers differs between *orb^ΔCPE^*and *orb^Δ3’UTR^*. In *orb^ΔCPE^*, Orb has a perinuclear and clamp-like localization in nurse cells of egg chambers, while in *orb^Δ3’UTR^* the Orb signal is much more diffuse.

The obtained mutations in the *orb* 3’UTR should not affect transcriptional regulation, although the results suggest that pre-mRNA levels are higher in *orb^Δ3’UTR^* and *orb^ΔCPE^* compared to WT. We suggest that these changes may be related to skewed proportions of somatic and germ cells or egg chambers at different stages of oogenesis in ovaries of mutant flies compared to control. This explanation seems more plausible though we can not rule out the possibility of direct changes in the transcriptional regulation of *orb* in mutant flies. While pre-mRNA levels differ between *orb^ΔCPE^*and *orb^Δ3’UTR^*, these mutants exhibit almost identical mRNA stability which is significantly lower than that of WT. This indicates that CPEs are responsible for *orb* mRNA stability.

The last 115 nucleotides of the *orb* 3’UTR are also essential for *orb* mRNA function, including its stability and translation activation. Removal of this sequence leads to more than a 90% decrease in *orb* mRNA stability in *orb^Δ3’UTR-DsRed^* and a decline of Orb protein to undetectable levels in ovaries of this line. This sequence contains nuclear polyadenylation signal but no CPE sites. The most probable cause for the decrease in *orb* mRNA stability is the removal of the polyadenylation signal, as the absence of a poly(A) tail triggers mRNA degradation. However, we found that an alternative polyadenylation signal appeared in the 3хPЗ promoter sequence, which is transcribed as part of the 3’UTR. Positive amplification of *orb* cDNA ends by the RACE procedure using an oligo(d)T primer confirms the presence of the *orb* mRNA poly(A) tail in *orb^Δ3’UTR-DsRed^*ovaries. We suggest that the 115-nt terminal sequence may contain additional *cis*-regulatory elements involved in maintaining the stability of *orb* mRNA molecules, apart from the PAS.

We consider *orb^Δ3’UTR-DsRed^* to be a null mutation. Similar morphological changes in ovaries were observed in two null mutations *orb^F343^* and *orb^dec^* in previous studies (Christerson and McKearin, 1994; Lantz et al., 1994; Rojas-Ríos et al., 2015). Our microscopy data suggest that in many germaria cysts fail to be encapsulated by follicular cells in region 2b or form cysts with an irregular number of cystocytes. The transition between regions 2a and 2b in the germarium is a checkpoint where defective germline cysts are eliminated by apoptosis (Drummond-Barbosa and Spradling, 2001). We assume that collapse of cysts in the germaria of *orb* null mutants is induced by their inability to pass the checkpoint in region 2a/2b. The structure of fusome appears normal in region 2a, but based on the presented figures, cysts with irregular numbers of cystocyes begin to form in region 2b when cysts are encapsulated by follicular cells. This issue requires further investigation. The effect of *orb* mutation on the increased number of cystocytes with round-shaped fusome in region 1 is an unexpected result, as Orb is not visualized in cystocytes at early stages of oogenesis. A recent study showed that both *orb* mRNA and protein are present in region 1 of the germarium, albeit at a minimal levels (Samuels et al., 2024).

Nurse cells are polyploid germ cells with polytene chromosomes undergoing 10-12 rounds of endocycles regulated by CyclinE/Cdk2 oscillations (Lilly and Duronio, 2005). Polytene chromosomes are visible as blob-like structures during the first four cycles. Starting from the fifth endocycle in Stage 6 egg chambers the cell cycles become truncated and late replicating heterochromatin is under-replicated. This leads to more dispersal chromatin organization, which is believed to promote ribosome synthesis and to be required for normal oocyte development (Dej and Spradling, 1999; Hammond and Laird, 1985; Volpe et al., 2001). Our microscopic analysis showed blob-like chromatin staining in nuclei of nurse cells in flies *orb^Δ3’UTR^*and *orb^ΔCPE^*. This staining pattern is visible along ovarioles in all egg chambers both containing only nurse cells and with one to three defective oocyte-like cells, which have weak Orb accumulation. This observation suggests that the *orb* 3’UTR mutations directly or indirectly affect the endoreplication in nurse cells. Hence, Orb may be essential to regulate specification and/or normal development of nurse cells, where Orb is presented in cytoplasm until late oogenesis (Costa et al., 2005; Lantz et al., 1994).

Nurse cells functioning may require Orb without activation of autoregulatory loop, which is inhibited in these cells (Wong and Schedl, 2011). Indeed, disruption of the Orb autoregulation brake oocyte development, while nurse cells are not significantly affected. In nurse cells, Orb may be required as both activator and repressor of translation, which depends on its phosphorylation status, like for other CPEB proteins. In *Drosophila* ovaries, Orb has been found as slow migrated hyperphosphorylated and fast migrated hypophosphorylated phosphoisoforms. These two phosphoisoforms are incorporated in different complexes separated by sucrose gradient fractionation. Hyperphosphorylated Orb is co-sedimented with 80S and polysome fractions, while hypophosphorylated Orb is co-sedimented with relatively small complexes enriched by translational repressors Bruno and Me31B (Wong et al., 2011). Observations of *orb* mutants give evidence that egg chamber formation cannot be successfully completed without both repressive and activating Orb phosphorisoforms. In *orb^F303^* ovaries, where only hypophosphorylated Orb has been detected, highly defective egg chambers are formed. The latter are without oocytes and without entire coating by follicle cell epithelium and their development stop soon thereafter formation (Lantz et al., 1994; Wong et al., 2011). In *orb^Δ3’UTR^* and *orb^ΔCPE^*, ovaries contain two Orb phosphoisoforms. In these both cases, egg chambers are formed without specified oocytes and develop until mid-oogenesis checkpoint stage.

Orb autoregulation is disrupted in *orb* 3’UTR mutants leading to failure of oocyte specification. At the same time, disruption of Orb autoregulation has no or little effect on nurse cell specification, while potentially causes abnormalities in their endoreplication. Based on these facts, we think that in ovaries Orb has two different functions. The first function is related with oocyte specification. It requires accumulation of high amount of Orb protein in one cyst cell and establishment of autoregulatory loop via interaction of Orb with its own mRNA. CPE motifs are essential cis-regulatory elements that provide *orb* mRNA accumulation and on-site translation. The second function is independent of Orb autoregulation and necessary for specification of nurse cells. Egg chambers are formed without oocyte and pass the checkpoint 2a/2b in the germarium even when Orb autoregulatory loop is disrupted. Germline cysts collapse at checkpoint 2a/2b if Orb protein is not expressed. Moreover, egg chamber formation requires both hyper- and hypophosphorylated Orb, which are translation activating and repressing forms of the protein.

## Materials and Methods

### Drosophila stocks

Flies were maintained on standard fly medium. To dissect ovaries, females were kept at 25° C and given fresh fly medium daily for 2-3 days. Ovaries were dissected from 2-5-day old females. For immunostaining flies were fed with yeast past on fly medium for 2-3 days. *w^1118^* flies were used as a wild type. Generation of *orb^Δ3’UTR^* by CRISPR/Cas9 system has been previously described (Barr et al., 2019b). *orb^resc^*was created by insertion of 24-1091nt fragment of 3’UTR into attP site in place of deletion in *orb^Δ3’UTR^*line using site-specific PhiC31 integrase. *orb^ΔCPE^* was generated by insertion of 30-1169 nt fragment of 3’UTR, which contains mutated CPE and CPE-like sequences. Sequences of the latter were given in an article of Stepien and co-authors (Stepien et al., 2016). *orb^Δ3’UTR-DsRed^*and *orb^ΔCPE-DsRed^* are intermediates of corresponding lines with *DsRed* marker. *orb^T840stop^* with a single point mutation was received by scarless CRISPR/Cas9 editing. *orb^915Δ3’UTR^* and *orb^915Δ3’UTR-DsRed^*are analogous to *orb^Δ3’UTR^*and *orb^Δ3’UTR-DsRed^*, but encode complete Orb protein.

### Western Blot

Ovaries were dissected and frozen at -80° C. Frozen ovaries were directly lysed and boiled in sample buffer (50 mM Tris-HCl, pH 6.8, 2% SDS, 10% glycerol, 100 mM DTT, 0.01% bromophenol blue) supplemented with 1×cOmplete Protease Inhibitor Cocktail (Roche, USA). To analyze impact of phosphorylation on Orb mobility in gel, 20 ovaries were lysed in 1% NP-40 and treated with 400 Units of λ protein phosphatase in PMP buffer supplemented with 1 mM MnCl2, 1× cOmplete Protease Inhibitor Cocktail and 0.8 mM PMSF for 30 min at 30° C (New England Biolabs, USA). Control samples were incubated in PNP buffer without MnCl2 and λ protein phosphatase and with 35 mM EDTA. Samples were resolved in 7.5% gel and transferred to 0.2µm PVDF membranes. Membranes were washed (PBS, 0.025% Tween 20) and blocked with 4% non-fat milk and incubated with mouse anti-Orb (4H8, concentrate, 1:1000) or anti-β-tubulin (E7, concentrate, 1:1000, Developmental Studies Hybridoma Bank, USA) antibody overnight at 4° C. Membranes were washed and incubated with HRP-conjugated goat anti-mouse antibodies (1:10000, Jackson Immunoresearch, USA) for 1 h. Then, membranes were washed and developed with Super Signal Western Femto substrate (ThermoFisher Scientific, USA). Chemiluminescence signal was detected by ChemiDoc visualization system (Bio-Rad, USA).

### Generation of polyclonal anti-Orb antibodies

Antibodies were generated against recombinant peptide corresponding to 410-560 amino acids of Orb, which does not include conserved domains (File S2). The fragment of *orb* was amplified from genomic DNA using primers with NdeI and XhoI restriction enzyme sites (Table S1). To generate histidine-tagged construct, XhoI/NdeI digested PCR-product was cloned into pET-28a treated by the same restriction enzymes. Peptide was expressed in BL21DE3 cells, purified on Ni-NTA agarose and used for immunization of rabbits. Antibodies were purified by affinity chromatography with antigen immobilized on Sepharose 4B columns. Specificity of the antibodies was tested on ovarian samples by Western Blot (Fig S8)

### Immunofluorescence

Ovaries were fixed with 4% PFA in PBS for 30 min. Ovaries were washed three times with PBST (0.1% Tween 20, PBS), permeabilized with PBXD (0.1% Tween 20, 0.3% Triton X-100, 0.3% sodium deoxycholate, PBS) for 1 h and washed again with PBST. The samples were treated with Image-iT FX Signal Enhancer (ThermoFisher Scientific, USA) and washed with PBST. Then, the samples were blocked with 5% normal goat serum (NGS) diluted with PBST for 1 h. The ovaries were incubated for 48 h at +4° C with primary antibodies diluted with 3% NGS in PBX (0.1% Tween 20, 0.3% Triton X-100, PBS). The following antibodies were used: rabbit polyclonal anti-Orb antibodies (440 ng/µl, see Methods above), rat anti-vasa antibody (supernatant, 1:15, anti-vasa, Developmental Studies Hybridoma Bank, USA), mouse anti-alpha-spectrin antibodies (concentrate, 1:180, 3A9, Developmental Studies Hybridoma Bank, USA). The ovaries were washed with PBST and incubated with goat anti-rabbit Alexa Fluor 488-conjugated antibodies (1:1000, ThermoFisher Scientific, USA) or donkey anti-rat Alexa Fluor 647-conjugated antibodies (1:200, Abcam, USA) and with phalloidin-Atto 633 (1:400, Sigma-Aldrich, USA) for 24 h at +4° C. After washing the samples were mounted in VectaShield with DAPI (Vector Laboratories, USA). Images were taken on a Stellaris 5 confocal microscope (Leica Microsystems, Germany) and primarily processed using Fiji software (Schindelin et al., 2012). Single optical sections were finally processed using Photoshop CS2 (Adobe, USA).

### RNA isolation, RT-qPCR and RACE

Dissected ovaries were frozen at -80° C. RNA isolation was performed by RNeasy Mini kit (Qiagen, USA). RNA concentration was determined by Qubit RNA HS kit (ThermoFisher Scientific, USA). The first strand of cDNA was synthetized by RNAscribe RT (Biolabmix, Russia) from 200 ng of total RNA and then diluted two times. To measure *orb* pre-mRNA, the samples were treated by DNase using DNA-free kit (ThermoFisher Scientific, USA). Then, cDNA was synthetized from 300 ng of treated RNA and diluted. RT-qPCR was conducted on CFX96 Touch Real-Time PCR Detection System (Bio-Rad, USA) using qPCRmix-HS SYBR master mix (Evrogen, Russia). 25-µl reaction mixture contained 1 μl (3 μl of *orb* pre-mRNA) of template cDNA and 0.25 μM of each primer with the following temperature program: 94°C for 30 s, 40 cycles of 94°C for 10 s, 56°C (58 °C for *orb* pre-mRNA) for 25 s and 72°C for 15 s. Each sample was analyzed in technical triplicate. In case of *orb* pre-mRNA measurement, the presence of genomic DNA was tested in DNase-treated RNA samples. Relative levels of expression were calculated by ΔΔCt method. *GAPDH2* and *Adh* were selected as reference genes because these genes are most stably expressed in ovaries with different mutations (personal communications). Levene’s test was applied to evaluate the equality of group variances. Statistical analysis was performed by one-way ANOVA with Fisher’s Least Significant Difference (LSD) test.

To determine sequences of the *orb* 3’UTR ends in females, we performed 3’-Step-Out RACE (Matz et al., 1999). Briefly, we generated cDNA using AAGCAGTGGTATCAACGCAGAGTAC(T)30VN primer (Evrogen, Russia). Then, each round of RACE was performed with gene-specific primer and primer mixture from Mint RACE primer set (Evrogen, Russia) using Encyclo polymerase of the same manufacturer. Primer sequences used in RT-qPCR and RACE are given in Table S1.

## Acknowledgements

We thank the Center for Precision Genome Editing and Genetic Technologies for Biomedicine, IGB RAS for the opportunity to work at Stellaris 5 confocal microscope (Leica Microsystems, Germany).

## References

Atkins, C. M., Davare, M. A., Oh, M. C., Derkach, V., and Soderling, T. R. (2005). Bidirectional regulation of cytoplasmic polyadenylation element-binding protein phosphorylation by Ca2+/calmodulin-dependent protein kinase II and protein phosphatase 1 during hippocampal long-term potentiation. J. Neurosci. 25, 5604–5610. doi:10.1523/JNEUROSCI.5051-04.2005.

Barr, J., Charania, S., Gilmutdinov, R., Yakovlev, K., Shidlovskii, Y., and Schedl, P. (2019a). The CPEB translational regulator, Orb, functions together with Par proteins to polarize the Drosophila oocyte. PLOS Genet. 15, e1008012. doi:10.1371/journal.pgen.1008012.

Barr, J., Diegmiller, R., Colonnetta, M. M., Ke, W., Imran Alsous, J., Stern, T., et al. (2024). To be or not to be: orb, the fusome and oocyte specification in Drosophila. Genetics 226. doi:10.1093/GENETICS/IYAE020.

Barr, J., Gilmutdinov, R., Wang, L., Shidlovskii, Y., and Schedl, P. (2019b). The Drosophila CPEB protein Orb specifies oocyte fate by a 3’ UTR-dependent autoregulatory loop. Genetics 213, 1431–1446. doi:10.1534/genetics.119.302687.

Benton, R., and St Johnston, D. (2003). Drosophila PAR-1 and 14-3-3 inhibit Bazooka/PAR-3 to establish complementary cortical domains in polarized cells. Cell 115, 691–704. doi:10.1016/S0092-8674(03)00938-3.

Bullock, S. L., Ringel, I., Ish-Horowicz, D., and Lukavsky, P. J. (2010). A’-form RNA helices are required for cytoplasmic mRNA transport in Drosophila. Nat. Struct. Mol. Biol. 17, 703–709. doi:10.1038/NSMB.1813.

Castagnetti, S., and Ephrussi, A. (2003). Orb and a long poly(A) tail are required for efficient oskar translation at the posterior pole of the Drosophila oocyte. Development 130, 835–843. doi:10.1242/DEV.00309.

Chang, J. S., Tan, L., and Schedl, P. (1999). The Drosophila CPEB homolog, orb, is required for oskar protein expression in oocytes. Dev. Biol. 215, 91–106. doi:10.1006/DBIO.1999.9444.

Chang, J. S., Tan, L., Wolf, M. R., and Schedl, P. (2001). Functioning of the Drosophila orb gene in gurken mRNA localization and translation. Development 128, 3169–3177. doi:10.1242/DEV.128.16.3169.

Charlesworth, A., Cox, L. L., and MacNicol, A. M. (2004). Cytoplasmic Polyadenylation Element (CPE)- and CPE-binding Protein (CPEB)-independent mechanisms regulate early class maternal mRNA translational activation in Xenopus oocytes. J. Biol. Chem. 279, 17650–17659. doi:10.1074/jbc.M313837200.

Charlesworth, A., Meijer, H. A., and De Moor, C. H. (2013). Specificity factors in cytoplasmic polyadenylation. Wiley Interdiscip. Rev. RNA 4, 437–461. doi:10.1002/wrna.1171.

Christerson, L. B., and McKearin, D. M. (1994). orb is required for anteroposterior and dorsoventral patterning during Drosophila oogenesis. Genes Dev. 8, 614–628. doi:10.1101/gad.8.5.614.

Costa, A., Wang, Y., Dockendorff, T. C., Erdjument-Bromage, H., Tempst, P., Schedl, P., et al. (2005). The Drosophila fragile X protein functions as a negative regulator in the orb autoregulatory pathway. Dev. Cell 8, 331–342. doi:10.1016/J.DEVCEL.2005.01.011.

Davidson, A., Parton, R. M., Rabouille, C., Weil, T. T., and Davis, I. (2016). Localized translation of gurken/TGF-α mRNA during axis specification is controlled by access to Orb/CPEB on processing bodies. Cell Rep. 14, 2451–2462. doi:10.1016/J.CELREP.2016.02.038.

De Cuevas, M., Lee, J. K., and Spradling, A. C. (1996). alpha-spectrin is required for germline cell division and differentiation in the Drosophila ovary. Development 122, 3959–3968. doi:10.1242/DEV.122.12.3959.

Dej, K. J., and Spradling, A. C. (1999). The endocycle controls nurse cell polytene chromosome structure during Drosophila oogenesis. Development 126, 293–303. doi:10.1242/DEV.126.2.293.

Drummond-Barbosa, D., and Spradling, A. C. (2001). Stem cells and their progeny respond to nutritional changes during Drosophila oogenesis. Dev. Biol. 231, 265–278. doi:10.1006/DBIO.2000.0135.

Duran-Arqué, B., Cañete, M., Castellazzi, C. L., Bartomeu, A., Ferrer-Caelles, A., Reina, O., et al. (2022). Comparative analyses of vertebrate CPEB proteins define two subfamilies with coordinated yet distinct functions in post-transcriptional gene regulation. Genome Biol. 23, 192-. doi:10.1186/S13059-022-02759-Y/FIGURES/5.

Findley, S. D., Tamanaha, M., Clegg, N. J., and Ruohola-Baker, H. (2003). Maelstrom, a Drosophila spindle-class gene, encodes a protein that colocalizes with Vasa and RDE1/AGO1 homolog, Aubergine, in nuage. Development 130, 859–871. doi:10.1242/DEV.00310.

Guillén-Boixet, J., Buzon, V., Salvatella, X., and Méndez, R. (2016). CPEB4 is regulated during cell cycle by ERK2/Cdk1-mediated phosphorylation and its assembly into liquid-like droplets. Elife 5. doi:10.7554/ELIFE.19298.

Hammond, M. P., and Laird, C. D. (1985). Chromosome structure and DNA replication in nurse and follicle cells of Drosophila melanogaster. Chromosoma 91, 267–278. doi:10.1007/BF00328222.

Hay, B., Jan, L. Y., and Jan, Y. N. (1988). A protein component of Drosophila polar granules is encoded by vasa and has extensive sequence similarity to ATP-dependent helicases. Cell 55, 577–587. doi:10.1016/0092-8674(88)90216-4.

Hodgman, R., Tay, J., Mendez, R., and Richter, J. D. (2001). CPEB phosphorylation and cytoplasmic polyadenylation are catalyzed by the kinase IAK1/Eg2 in maturing mouse oocytes. Development 128, 2815–2822. doi:10.1242/DEV.128.14.2815.

Huang, Y. S., Jung, M. Y., Sarkissian, M., and Richter, J. D. (2002). N-methyl-D-aspartate receptor signaling results in Aurora kinase-catalyzed CPEB phosphorylation and alpha CaMKII mRNA polyadenylation at synapses. EMBO J. 21, 2139–2148. doi:10.1093/EMBOJ/21.9.2139.

Huynh, J. R., and St Johnston, D. (2000). The role of BicD, Egl, Orb and the microtubules in the restriction of meiosis to the Drosophila oocyte. Development 127, 2785–2794. doi:10.1242/DEV.127.13.2785.

Ivshina, M., Lasko, P., and Richter, J. D. (2014). Cytoplasmic polyadenylation element binding proteins in development, health, and disease. Annu. Rev. Cell Dev. Biol. 30, 393–415. doi:10.1146/ANNUREV-CELLBIO-101011-155831.

Katsioudi, G., Dreos, R., Arpa, E. S., Gaspari, S., Liechti, A., Sato, M., et al. (2023). A conditional Smg6 mutant mouse model reveals circadian clock regulation through the nonsense-mediated mRNA decay pathway. Sci. Adv. 9. doi:10.1126/SCIADV.ADE2828.

Kortenoeven, M. L. A., Schweer, H., Cox, R., Wetzels, J. F. M., and Deen, P. M. T. (2012). Lithium reduces aquaporin-2 transcription independent of prostaglandins. Am. J. Physiol. Cell Physiol. 302. doi:10.1152/AJPCELL.00197.2011.

Kozlov, E., Shidlovskii, Y. V., Gilmutdinov, R., Schedl, P., and Zhukova, M. (2021). The role of CPEB family proteins in the nervous system function in the norm and pathology. Cell Biosci. 11. doi:10.1186/S13578-021-00577-6.

Lantz, V., Ambrosio, L., and Schedl, P. (1992). The Drosophila orb gene is predicted to encode sex-specific germline RNA-binding proteins and has localized transcripts in ovaries and early embryos | Development. Development 115, 75–88. Available at: https://dev.biologists.org/content/115/1/75.long [Accessed June 27, 2020].

Lantz, V., Chang, J. S., Horabin, J. I., Bopp, D., and Schedl, P. (1994). The Drosophila orb RNA-binding protein is required for the formation of the egg chamber and establishment of polarity. Genes Dev. 8, 598–613. doi:10.1101/gad.8.5.598.

Lantz, V., and Schedl, P. (1994). Multiple cis-acting targeting sequences are required for orb mRNA localization during Drosophila oogenesis. Mol. Cell. Biol. 14, 2235–42. doi:10.1128/mcb.14.4.2235.

Lasko, P. F., and Ashburner, M. (1988). The product of the Drosophila gene vasa is very similar to eukaryotic initiation factor-4A. Nature 335, 611–617. doi:10.1038/335611A0.

Lasko, P. F., and Ashburner, M. (1990). Posterior localization of vasa protein correlates with, but is not sufficient for, pole cell development. Genes Dev. 4, 905–921. doi:10.1101/GAD.4.6.905.

Lee, J. K., Brandin, E., Branton, D., and Goldstein, L. S. B. (1997). alpha-Spectrin is required for ovarian follicle monolayer integrity in Drosophila melanogaster. Development 124, 353–362. doi:10.1242/DEV.124.2.353.

Lilly, M. A., and Duronio, R. J. (2005). New insights into cell cycle control from the Drosophila endocycle. Oncogene 24, 2765–2775. doi:10.1038/SJ.ONC.1208610.

Lin, H., Yue, L., and Spradling, A. C. (1994). The Drosophila fusome, a germline-specific organelle, contains membrane skeletal proteins and functions in cyst formation. Development 120, 947–956. doi:10.1242/DEV.120.4.947.

McCall, K. (2004). Eggs over easy: cell death in the Drosophila ovary. Dev. Biol. 274, 3–14. doi:10.1016/J.YDBIO.2004.07.017.

Mendez, R., Hake, L. E., Andresson, T., Littlepage, L. E., Ruderman, J. V., and Richter, J. D. (2000a). Phosphorylation of CPE binding factor by Eg2 regulates translation of c-mos mRNA. Nature 404, 302–307. doi:10.1038/35005126.

Mendez, R., Murthy, K. G. K., Ryan, K., Manley, J. L., and Richter, J. D. (2000b). Phosphorylation of CPEB by Eg2 mediates the recruitment of CPSF into an active cytoplasmic polyadenylation complex. Mol. Cell 6, 1253–1259. doi:10.1016/S1097-2765(00)00121-0.

Nezis, I. P., Stravopodis, D. J., Papassideri, I., Robert-Nicoud, M., and Margaritis, L. H. (2000). Stage-specific apoptotic patterns during Drosophila oogenesis. Eur. J. Cell Biol. 79, 610–620. doi:10.1078/0171-9335-00088.

Pai, T. P., Chen, C. C., Lin, H. H., Chin, A. L., Lai, J. S. Y., Lee, P. T., et al. (2013). Drosophila Orb protein in two mushroom body output neurons is necessary for long-term memory formation. Proc. Natl. Acad. Sci. U. S. A. 110, 7898–7903. doi:10.1073/pnas.1216336110.

Peterson, J. S., Barkett, M., and McCall, K. (2003). Stage-specific regulation of caspase activity in Drosophila oogenesis. Dev. Biol. 260, 113–123. doi:10.1016/S0012-1606(03)00240-9.

Retelska, D., Iseli, C., Bucher, P., Jongeneel, C. V., and Naef, F. (2006). Similarities and differences of polyadenylation signals in human and fly. BMC Genomics 7. doi:10.1186/1471-2164-7-176.

Rojas-Ríos, P., Chartier, A., Pierson, S., Séverac, D., Dantec, C., Busseau, I., et al. (2015). Translational control of autophagy by Orb in the Drosophila germline. Dev. Cell 35, 622–631. doi:10.1016/J.DEVCEL.2015.11.003.

Samuels, T. J., Gui, J., Gebert, D., and Karam Teixeira, F. (2024). Two distinct waves of transcriptome and translatome changes drive Drosophila germline stem cell differentiation. EMBO J. 43, 1591. doi:10.1038/S44318-024-00070-Z.

Sanfilippo, P., Wen, J., and Lai, E. C. (2017). Landscape and evolution of tissue-specific alternative polyadenylation across Drosophila species. Genome Biol. 18. doi:10.1186/S13059-017-1358-0.

Smibert, P., Miura, P., Westholm, J. O., Shenker, S., May, G., Duff, M. O., et al. (2012). Global patterns of tissue-specific alternative polyadenylation in Drosophila. Cell Rep. 1, 277–289. doi:10.1016/J.CELREP.2012.01.001.

Stepien, B. K., Oppitz, C., Gerlach, D., Dag, U., Novatchkova, M., Krüttner, S., et al. (2016). RNA-binding profiles of Drosophila CPEB proteins Orb and Orb2. Proc. Natl. Acad. Sci. 113, E7030–E7038. doi:10.1073/pnas.1603715113.

Styhler, S., Nakamura, A., Swan, A., Suter, B., and Lasko, P. (1998). vasa is required for GURKEN accumulation in the oocyte, and is involved in oocyte differentiation and germline cyst development. Development 125, 1569–1578. doi:10.1242/DEV.125.9.1569.

Tan, L., Chang, J. S., Costa, A., and Schedl, P. (2001). An autoregulatory feedback loop directs the localized expression of the Drosophila CPEB protein Orb in the developing oocyte. Development 128, 1159–1169. doi:10.1242/DEV.128.7.1159.

Tay, J., Hodgman, R., and Richter, J. D. (2000). The control of cyclin B1 mRNA translation during mouse oocyte maturation. Dev. Biol. 221, 1–9. doi:10.1006/DBIO.2000.9669.

Tian, B., and Graber, J. H. (2012). Signals for pre-mRNA cleavage and polyadenylation. Wiley Interdiscip. Rev. RNA 3, 385. doi:10.1002/WRNA.116.

Volpe, A. M., Horowitz, H., Grafer, C. M., Jackson, S. M., and Berg, C. A. (2001). Drosophila rhino encodes a female-specific chromo-domain protein that affects chromosome structure and egg polarity. Genetics 159, 1117–1134. doi:10.1093/GENETICS/159.3.1117.

Wong, L. C., Costa, A., McLeod, I., Sarkeshik, A., Yates, J., Kyin, S., et al. (2011). The functioning of the Drosophila CPEB protein Orb is regulated by phosphorylation and requires casein kinase 2 activity. PLoS One 6, e24355. doi:10.1371/JOURNAL.PONE.0024355.

Wong, L. C., and Schedl, P. (2011). Cup blocks the precocious activation of the Orb autoregulatory loop. PLoS One 6. doi:10.1371/journal.pone.0028261.

