## Supplementary material for "Stability of *orb* mRNA in *Drosophila* ovaries is regulated by *cis*-regulatory elements in the 3’ UTR and influences germ cell specification": Figures S1-S8

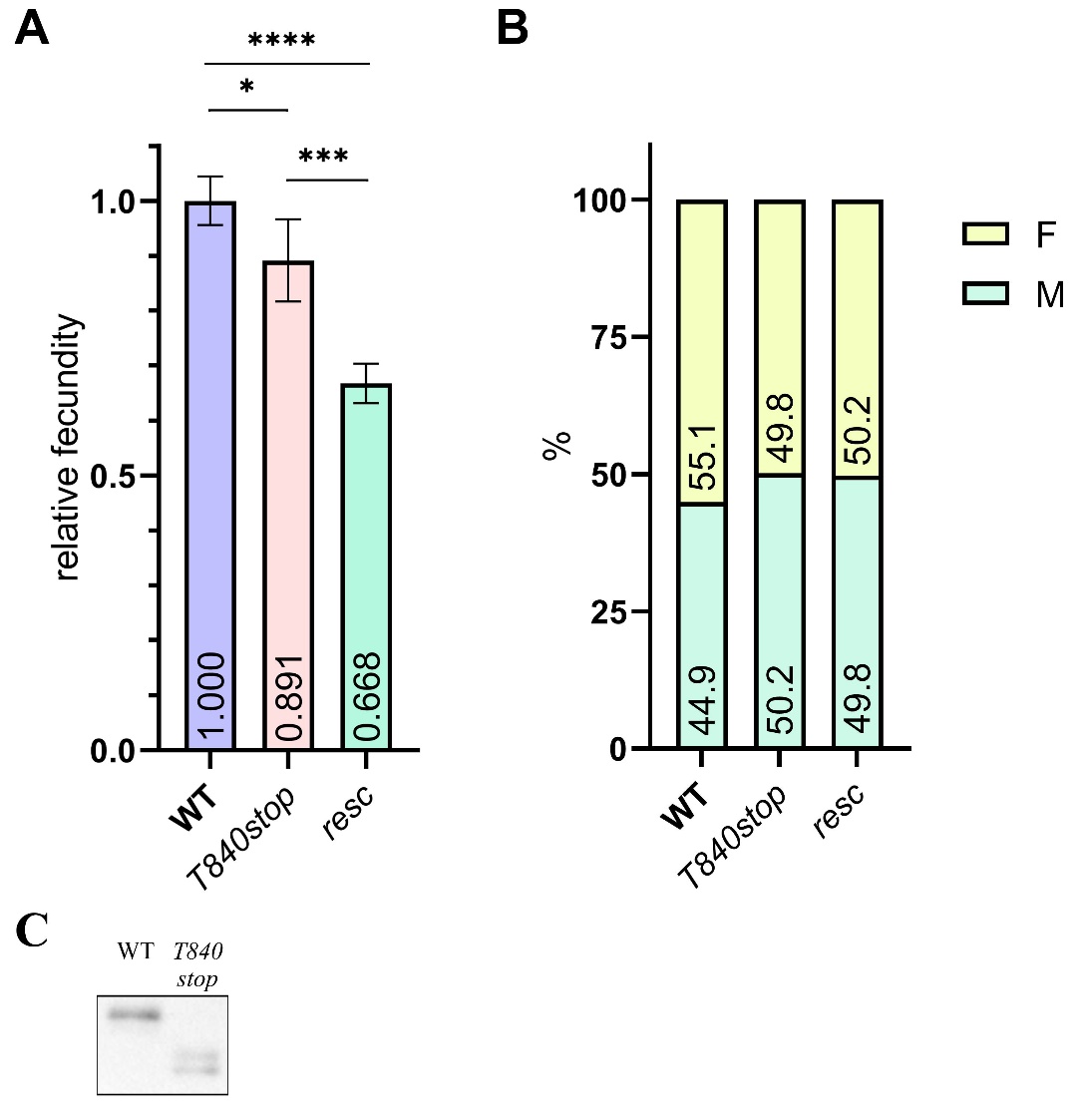

**Fig S1. Fecundity and sex ratio in wild type (WT), *orb^T840stop^* and *orb^resc^* homozygotes.** 1-2-d flies (10 females and 10 males per tube) were kept at 25° C and transferred to fresh tubes daily for two days. After then, the flies were kept in tubes with fresh food for egg laying for three days. Adult flies were counted after week from first hatching. (**A**) Relative fecundity of the mutants was calculated in relation to the WT. Values are mean±SD (n = 4). Statistical significance was counted using a one-way ANOVA with subsequent Tukey post-hoc test (* p < 0.05, *** p < 0.001, **** p < 0.0001). (**B**) Sex ratio among hatched flies. WT (n = 608), *orb^T840stop^* (n = 542) *orb^resc^* (n = 406). (**C**) Western Blot analysis shows expression of shortened Orb in the *orb^T840stop^* ovaries.

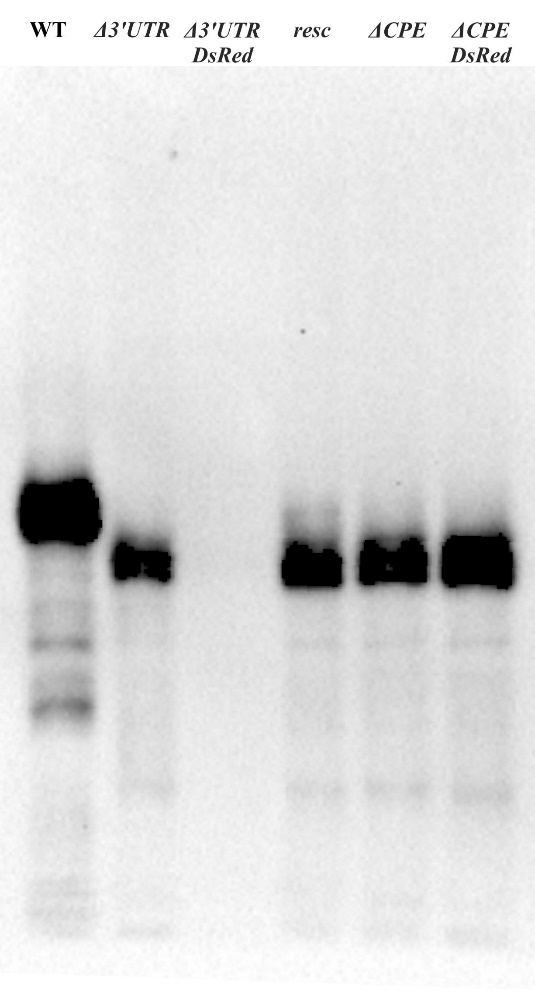

**Fig S2.** Western blot analysis of Orb in 3-5-day ovaries using 4H8 antibody. Membrane overexposure is to detect potential minimal Orb amount in *orb^Δ3’UTR-DsRed^*.

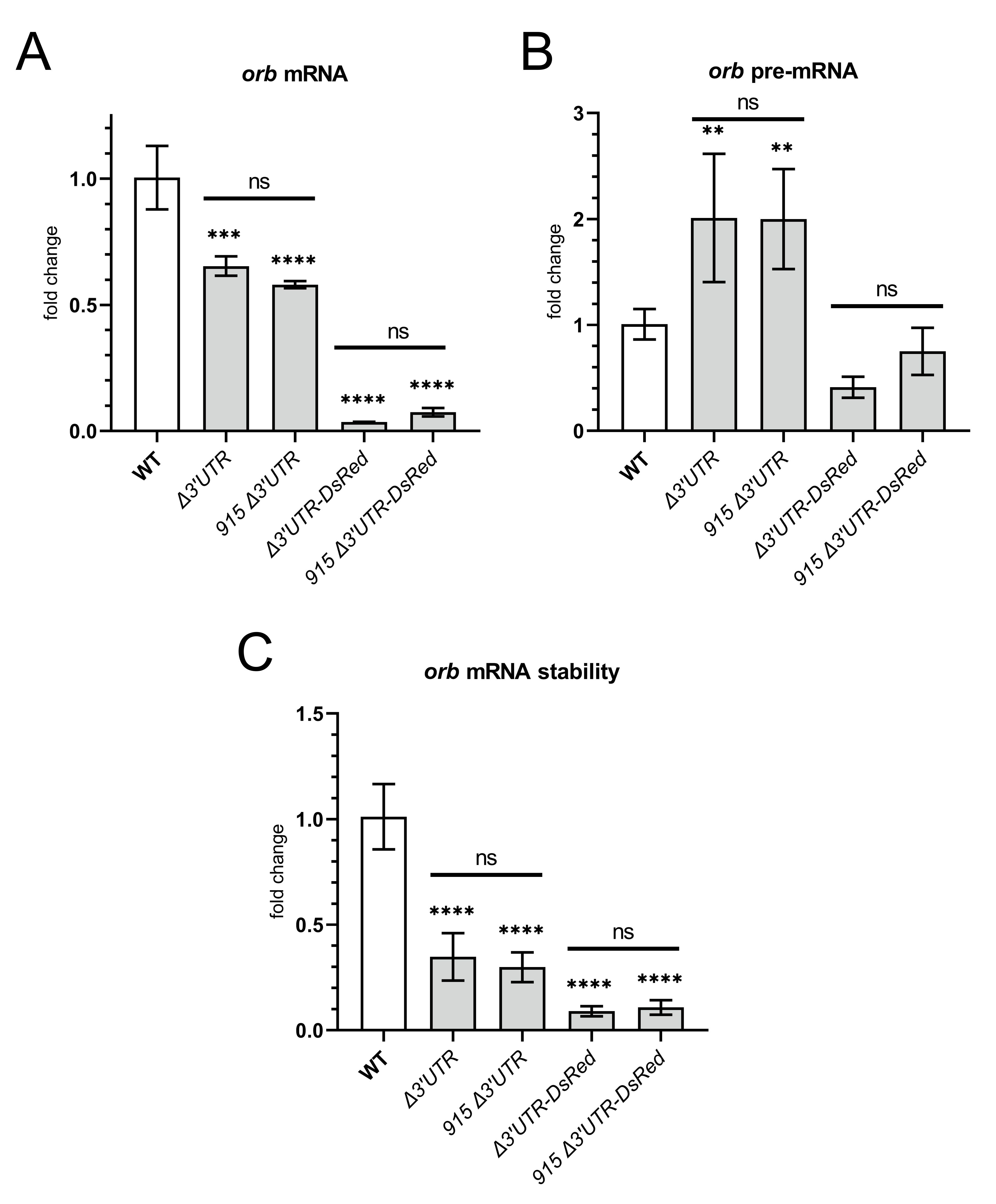

**Fig S3.** Expression of *orb* in ovaries of *orb^Δ3’UTR^* and *orb^Δ3’UTR-DsRed^* females estimated by RT-qPCR. (**A**) Expression of *orb* mRNA (WT and *orb^Δ3’UTR^* n=2, *orb^915 Δ3’UTR^* and *orb^915 Δ3’UTR-DsRed^* n=3). (**B**) Evaluation of the *orb* transcriptional levels by measurement of *orb* pre-mRNA (n=3). (**C**) Levels of the *orb* mRNA stability calculated by ratio of relative level of mRNA to relative level of pre-mRNA. Values are shown as mean ± SD. Statistical significance estimated by comparison test with WT is shown. ** p < 0.01, *** p < 0.001, **** p < 0.0001. “ns” above lines mean non-significant statistical differences between columns.

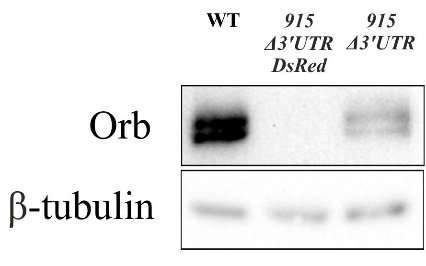

**Fig S4.** Western blot analysis of the *orb^915 Δ3’UTR^* and *orb^915 Δ3’UTR-DsRed^* ovaries expressed full-size Orb protein. β-tubulin was used as loading control. No Orb signal was detected in

*orb^915 Δ3’UTR-DsRed^* ovaries.

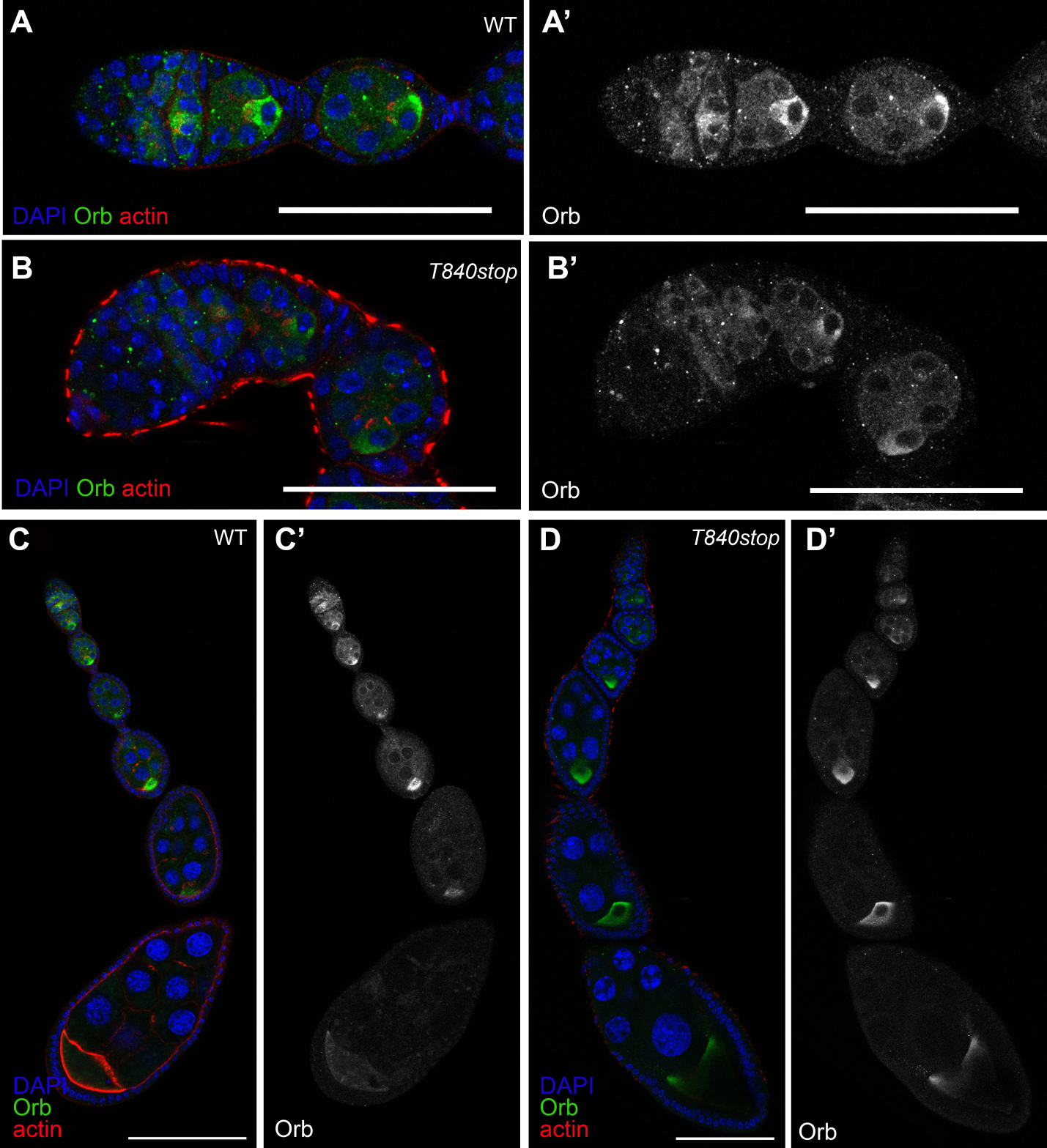

**Figure S5.** Orb protein localization in ovaries of WT and *orb^T840stop^*. Ovaries are stained with antibodies against Orb protein (green), DAPI (blue) and Alexa Fluor 633 phalloidin (red). Panels with apostrophes (**A’-D’**) demonstrate separately a channel with Orb staining. (**A, B**) – Orb protein in germaria; (**C, D**) **–** Orb has similar localization in ovarioles of of WT and *orb^T840stop^.* Bars are 50 µm (A-B’), 100 µm (C-D’).

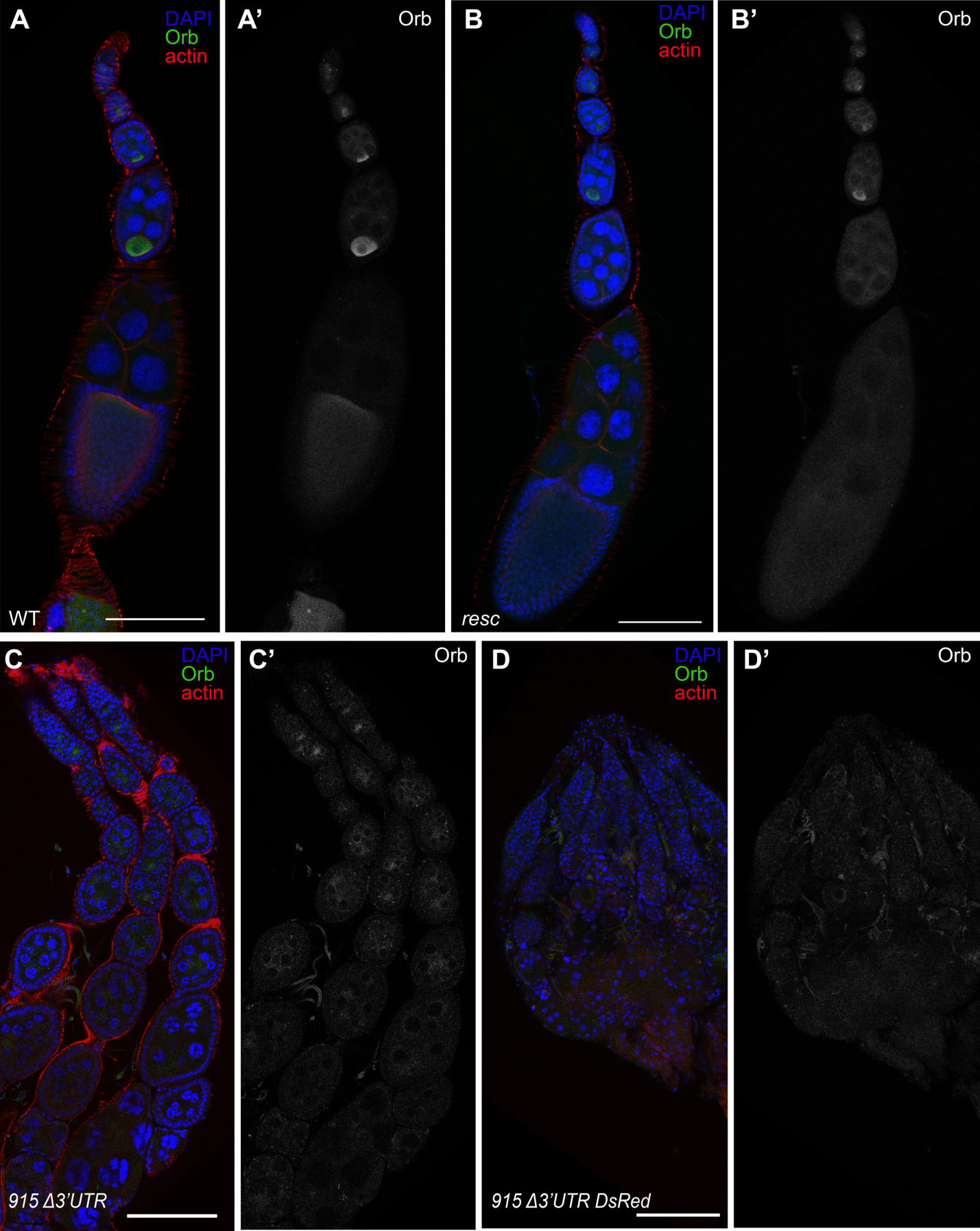

**Fig S6**. Orb localization in ovaries of different mutant flies. Ovaries are stained with antibodies against Orb protein (green), DAPI (blue) and Alexa Fluor 633 phalloidin (red). Separate images of channel with Orb staining are presented on the right (**A’-D’**). In WT (**A**, **A’**) and *orb^resc^* (**B**, **B’**) Orb is localized similarly, it is specific to oocytes-specific in egg chambers. In *orb^915Δ3’UTR^* (**C**, **C’**), oocytes are absent and a weak diffuse Orb signal localizes in cytoplasm of nurse cells, similar to *orb^Δ3’UTR^*. In *orb^915Δ3’UTR-DsRed^* (**D**, **D’**), egg chambers are not visible. Ovarioles are thin and without Orb signal, similar to *orb^Δ3’UTR-DsRed^*. Bars are 100 µm.

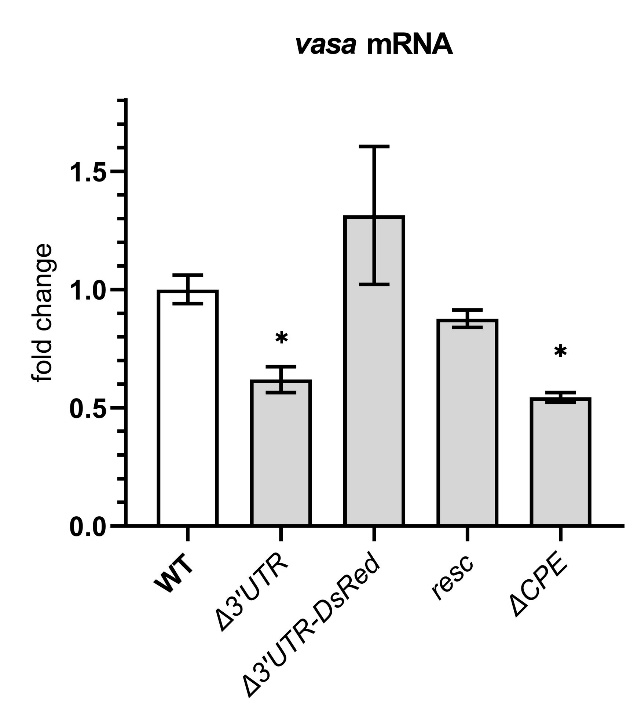

**Fig S7.** Expression of *vasa* in ovaries estimated by RT-qPCR. Values are shown as mean ± SD (Standard deviation) (n=2). Statistical significance estimated by comparison test with WT is shown. * p < 0.05.

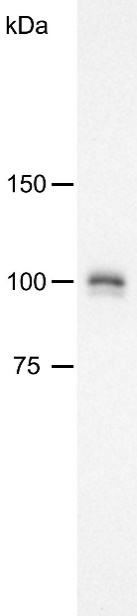

**Fig S8.** Analysis of specificity of rabbit polyclonal anti-Orb antibodies. Antibodies were used in concentration of 114 ng/ml. As seen in the Figure, Orb is detected as two bands with approximate weight of 100 kDa in WT ovarian lysate.
