## Supplementary material for "Stability of *orb* mRNA in *Drosophila* ovaries is regulated by *cis*-regulatory elements in the 3’ UTR and influences germ cell specification": File S1

**Sequences of 3’UTRs of the *orb* mRNA**

Consensus CPE are marked purple.

G-containing CPE are marked green.

Sequences similar to CPEs are marked grey.

In *orb^ΔCPE^* and *orb^ΔCPE-DsRed+^*, sequences, where CPE and CPE-related sequences had been deleted are marked yellow.

PAS sequences close to the 3’UTR ends are underlined.

WT (w^1118^)

It includes the 3’ end of the translated region

AACGAAGTATCAGGGACAACAGAATGGGGAACGGTCAGCATCAACAGCACCAGCAGCATCAGTACCAACAGCAGAAACACCGTCAGCTGCAAGAATACTCGCAGCCCCACAGTCTCAACGTGATGGGAAACTCAGGAGCTGCCAATGCCGCTGCTACATCAATGGTAACTCTTCAGCAGCGGCAAATTCACAAGGTGCGAATACAGCGTCAACAGCATCAGGCGATCTAGACTGTTACGGCTTTTATCCACACCGTTTTAACGGATGTTTCCGCAATATAATGTGTGAGACTTTGGACTTGTAGGCGACGTAGAAATATGGCGGAGATAGTTTTCTTGCAGCCGATGGAGACCCGGGCTCATCATCGTCATTATCATTGTAAACACAGTTTTATTGATGTGTATATAATATCGGTACAATATGAAGCTTACACTTTGAGTTTCATTTAAGATAATTATTGTTAGACGACTCCCCCAAAAACGAACACCCTCGAATCGAAACAGAAGAGAGACCCCCATTCATACTATTTTTTGATAATCATGCTTACATATCAGCATTTGGAGCTGGCATTCGAATGCTAAATGAATGATACATGAAATGAGCAATTTCATTTACTCATACCCCTCATACAAATACTACTCAAGTTACTTTTTGGCTAAGAAAGCAGTTATTAATTTCTAATTTGTATGTTTAATTTTAAGGAAACGAAACCGCGCGGCAAATGGCTACTTGAAATGTCTCGACCAATTTGCCGCGCTGCGAGATTAGAAACACCATTTTTGTTATGTTTTGTGCTATTACTGAGGAATTTTATGTAGTTTCTTTTTACATAGCCAAGCCCCCGACTCGAGTTAATATGATATTATATTATTTGGATTGTCCGCTAAGCGTTTATCAGGAATTTCAATTTTTAAGAAAACATTTTAAAAATTGTAAATTCGTTTAACTCACCAGTCTCCCTTTTTTGTTTTTCATTGAATTGTTACACACATGAGTTATTATAAATATACTGTTTAGTTTATTTTTTGTTTTGGATTAGCTTTAAGCATTATCCTTGTGAACATTAACGCGATGCCTGATTGATTGTTGAACTATGTTTTAAGCATTTATCATTCTTTGGCTTTTTTGGCTGTTCCAGTTTTATAGCTCATGGGCAATAAGCATATAATTTATGTCTACTTTATTTTTCATGTATTTTGAGAAAGGATCTCTCTAGCGCTTATCTATAGAAAACACACATATGTATGTATACAAAAATATTACATAGTGTACATTACAAGGCTTTATATAATTTAAAACTTGATAAGTTTGTAAACCTACACAGAATAGGAAAAAAATACTTAAGTCTATATCTTAAACACGAATTGATCACTAAACGATAAAATAACGCACACACACCTATTCACCAACCAAACAAATATACAAAATCGCTATTG

*orb^resc^*

AATGAAGTATCAGGGACAACAGAATGGGGAACGGTCAGCATCAACAGCACCAGCAGCATCAGTACCAACAGCAGAAACACCGTCAGCTGCAAGAATACTCGCAGCCCCACAGTCTCAACGTGATGGGAAACTCAGGAGCTGCCAATGCCGCTGCTACATCAATGGTAACTCTTCAGCAGCGGCAAATTCACAAGGTGCGAATACAGCGTCAACAGCATCAGGCGATCTAGACTGTTACGGCTTTTATCCACACGCGGCCGCGGACATATGCACACCTGCGATCGTAGTGCCCCAACTGGGGTAACCTTTGGGCTCCCCGGGCGCGTACTCCACCTCGAGTCTAGAGAATTCGGATCCGTTTTAACGGATGTTTCCGCAATATAATGTGTGAGACTTTGGACTTGTAGGCGACGTAGAAATATGGCGGAGATAGTTTTCTTGCAGCCGATGGAGACCCGGGCTCATCATCGTCATTATCATTGTAAACACAGTTTTATTGATGTGTATATAATATCGGTACAATATGAAGCTTACACTTTGAGTTTCATTTAAGATAATTATTGTTAGACGACTCCCCCAAAAACGAACACCCTCGAATCGAAACAGAAGAGAGACCCCCATTCATACTATTTTTTGATAATCATGCTTACATATCAGCATTTGGAGCTGGCATTCGAATGCTAAATGAATGATACATGAAATGAGCAATTTCATTTACTCATACCCCTCATACAAATACTACTCAAGTTACTTTTTGGCTAAGAAAGCAGTTATTAATTTCTAATTTGTATGTTTAATTTTAAGGAAACGAAACCGCGCGGCAAATGGCTACTTGAAATGTCTCGACCAATTTGCCGCGCTGCGAGATTAGAAACACCATTTTTGTTATGTTTTGTGCTATTACTGAGGAATTTTATGTAGTTTCTTTTTACATAGCCAAGCCCCCGACTCGAGTTAATATGATATTATATTATTTGGATTGTCCGCTAAGCGTTTATCAGGAATTTCAATTTTTAAGAAAACATTTTAAAAATTGTAAATTCGTTTAACTCACCAGTCTCCCTTTTTTGTTTTTCATTGAATTGTTACACACATGAGTTATTATAAATATACTGTTTAGTTTATTTTTTGTTTTGGATTAGCTTTAAGCATTATCCTTGTGAACATTAACGCGATGCCTGATTGATTGTTGAACTATGTTTTAAGCATTTATCATTCTTTGGCTTTTTTGGCTGTTCCAGTTTTATAGCTCATGGGCAATAAGCATATAATTTATGTCTACTTTATTTTTCATGTATTTTGAGAAAGGATCTCTCTAGCGCTTATCTATAGAAAACACACATATGTATGTATACAAAAATATTACATAGTGTACATTACAAGGCTTTATATAATTTAAAACTTGATAAGTTTGTAAACCTACAGATCTCGTACGATAACTTCGTATAATGTATGCTATACGAAGTTATAGAAGAGCACTAGTAAAGATCTCAGAATAGGAAAAAAATACTTAAGTCTATATCTTAAACACGAATTGATCACTAAACGATAAAATAACGCACACACACCTATTCACCAACCAAACAAATATACAAAATCGCTATTG

*orb^ΔCPE^*

AATGAAGTATCAGGGACAACAGAATGGGGAACGGTCAGCATCAACAGCACCAGCAGCATCAGTACCAACAGCAGAAACACCGTCAGCTGCAAGAATACTCGCAGCCCCACAGTCTCAACGTGATGGGAAACTCAGGAGCTGCCAATGCCGCTGCTACATCAATGGTAACTCTTCAGCAGCGGCAAATTCACAAGGTGCGAATACAGCGTCAACAGCATCAGGCGATCTAGACTGTTACGGCTTTTATCCACACGCGGCCGCGGACATATGCACACCTGCGATCGTAGTGCCCCAACTGGGGTAACCTTTGGGCTCCCCGGGCGCGTACTCCACCTCGAGTCTAGAGAATTCCGTAACGGATGTTTCCGCAATATAATGTGTGAGACTTTGGACTTGTAGGCGACGTAGAAATATGGCGGAGATAGTTTTCTTGCAGCCGATGGAGACCCGGGCTCATCATCGTCATTATCATTGTAAACACAGTATTGATGTGTATATAATATCGGTACAATATGAAGCTTACACTTTGAGTTTCATTTAAGATAATTATTGTTAGACGACTCCCCCAAAAACGAACACCCTCGAATCGAAACAGAAGAGAGACCCCCATTCATACTATTTTTTGATAATCATGCTTACATATCAGCATTTGGAGCTGGCATTCGAATGCTAAATGAATGATACATGAAATGAGCAATTTCATTTACTCATACCCCTCATACAAATACTACTCAAGTTACTTTTTGGCTAAGAAAGCAGTTATTAATTTCTAATTTGTATGTTTAATAAGGAAACGAAACCGCGCGGCAAATGGCTACTTGAAATGTCTCGACCAATTTGCCGCGCTGCGAGATTAGAAACACCATGTTATGTGTGCTATTACTGAGGAATATGTAGTTTCTACATAGCCAAGCCCCCGACTCGAGTTAATATGATATTATATTATTTGGATTGTCCGCTAAGCGTTTATCAGGAATTTCAATAAGAAAACATAAAAATTGTAAATTCGTTTAACTCACCAGTCTCCCTTGTTTTTCATTGAATTGTTACACACATGAGTTATTATAAATATACTGTTTAGTTTATTGTTTTGGATTAGCTTTAAGCATTATCCTTGTGAACATTAACGCGATGCCTGATTGATTGTTGAACTATGTAAGCATTTATCATTCTTTGGCTTTTTTGGCTGTTCCAGTATAGCTCATGGGCAATAAGCATATAATTTATGTCTACTTTATTTTTCATGTATTTTGAGAAAGGATCTCTCTAGCGCTTATCTATAGAAAACACACATATGTATGTATACAAAAATATTACATAGTGTACATTACAAGGCTTTATATAATTTAAAACTTGATAAGTTTGTAAACCTACACAGAATAGGAAAAAAATACTTAAGTCTATATCTTAAACACGAATTGATCACTAAACGATAAAATAACGCACACACACCCGTACGATAACTTCGTATAATGTATGCTATACGAAGTTATAGAAGAGCACTAGTAAAGATCTCAGAATAGGAAAAAAATACTTAAGTCTATATCTTAAACACGAATTGATCACTAAACGATAAAATAACGCACACACACCTATTCACCAACCAAACAAATATACAAAATCGCTATTG

*orb^ΔCPE-DsRed^*

AATGAAGTATCAGGGACAACAGAATGGGGAACGGTCAGCATCAACAGCACCAGCAGCATCAGTACCAACAGCAGAAACACCGTCAGCTGCAAGAATACTCGCAGCCCCACAGTCTCAACGTGATGGGAAACTCAGGAGCTGCCAATGCCGCTGCTACATCAATGGTAACTCTTCAGCAGCGGCAAATTCACAAGGTGCGAATACAGCGTCAACAGCATCAGGCGATCTAGACTGTTACGGCTTTTATCCACACGCGGCCGCGGACATATGCACACCTGCGATCGTAGTGCCCCAACTGGGGTAACCTTTGGGCTCCCCGGGCGCGTACTCCACCTCGAGTCTAGAGAATTCCGTAACGGATGTTTCCGCAATATAATGTGTGAGACTTTGGACTTGTAGGCGACGTAGAAATATGGCGGAGATAGTTTTCTTGCAGCCGATGGAGACCCGGGCTCATCATCGTCATTATCATTGTAAACACAGTATTGATGTGTATATAATATCGGTACAATATGAAGCTTACACTTTGAGTTTCATTTAAGATAATTATTGTTAGACGACTCCCCCAAAAACGAACACCCTCGAATCGAAACAGAAGAGAGACCCCCATTCATACTATTTTTTGATAATCATGCTTACATATCAGCATTTGGAGCTGGCATTCGAATGCTAAATGAATGATACATGAAATGAGCAATTTCATTTACTCATACCCCTCATACAAATACTACTCAAGTTACTTTTTGGCTAAGAAAGCAGTTATTAATTTCTAATTTGTATGTTTAATAAGGAAACGAAACCGCGCGGCAAATGGCTACTTGAAATGTCTCGACCAATTTGCCGCGCTGCGAGATTAGAAACACCATGTTATGTGTGCTATTACTGAGGAATATGTAGTTTCTACATAGCCAAGCCCCCGACTCGAGTTAATATGATATTATATTATTTGGATTGTCCGCTAAGCGTTTATCAGGAATTTCAATAAGAAAACATAAAAATTGTAAATTCGTTTAACTCACCAGTCTCCCTTGTTTTTCATTGAATTGTTACACACATGAGTTATTATAAATATACTGTTTAGTTTATTGTTTTGGATTAGCTTTAAGCATTATCCTTGTGAACATTAACGCGATGCCTGATTGATTGTTGAACTATGTAAGCATTTATCATTCTTTGGCTTTTTTGGCTGTTCCAGTATAGCTCATGGGCAATAAGCATATAATTTATGTCTACTTTATTTTTCATGTATTTTGAGAAAGGATCTCTCTAGCGCTTATCTATAGAAAACACACATATGTATGTATACAAAAATATTACATAGTGTACATTACAAGGCTTTATATAATTTAAAACTTGATAAGTTTGTAAACCTACACAGAATAGGAAAAAAATACTTAAGTCTATATCTTAAACACGAATTGATCACTAAACGATAAAATAACGCACACACACCCGTACGATAACTTCGTATAATGTATGCTATACGAAGTTATGTCGACGAATTGCTAGCTGGCGGCCGCACCGGTTAAGATACATTGATGAGTTTGGACAAACCACAACTAGAATGCAGTGAAAAAAATGCTTTATTTGTGAAATTTGTGATGCTATTGCTTTATTTGTAACCATTATAAGCTGCAATAAACAAGTTAACAAC

*orb^Δ3’UTR^*

AATGAAGTATCAGGGACAACAGAATGGGGAACGGTCAGCATCAACAGCACCAGCAGCATCAGTACCAACAGCAGAAACACCGTCAGCTGCAAGAATACTCGCAGCCCCACAGTCTCAACGTGATGGGAAACTCAGGAGCTGCCAATGCCGCTGCTACATCAATGGTAACTCTTCAGCAGCGGCAAATTCACAAGGTGCGAATACAGCGTCAACAGCATCAGGCGATCTAGACTGTTACGGCTTTTATCCACACGCGGCCGCGGACATATGCACACCTGCGATCGTAGTGCCCCAACTGGGGTAACCTTTGAGTTCTCTCAGTTGGGGGCGTAGATAACTTCGTATAATGTATGCTATACGAAGTTATAGAAGAGCACTAGTAAAGATCTCAGAATAGGAAAAAAATACTTAAGTCTATATCTTAAACACGAATTGATCACTAAACGATAAAATAACGCACACACACCTATTCACCAACCAAACAAATATACAAAATCGCTATTG

*orb^Δ3’UTR-DsRed^*

AATGAAGTATCAGGGACAACAGAATGGGGAACGGTCAGCATCAACAGCACCAGCAGCATCAGTACCAACAGCAGAAACACCGTCAGCTGCAAGAATACTCGCAGCCCCACAGTCTCAACGTGATGGGAAACTCAGGAGCTGCCAATGCCGCTGCTACATCAATGGTAACTCTTCAGCAGCGGCAAATTCACAAGGTGCGAATACAGCGTCAACAGCATCAGGCGATCTAGACTGTTACGGCTTTTATCCACACGCGGCCGCGGACATATGCACACCTGCGATCGTAGTGCCCCAACTGGGGTAACCTTTGAGTTCTCTCAGTTGGGGGCGTAGATAACTTCGTATAATGTATGCTATACGAAGTTATCGTACGGGATCTAATTCAATTAGAGACTAATTCAATTAGAGCTAATTCAATTAGGATCCAAGCTTATCGATTTCGAACCCTCGACCGCCGGAGTATAAATAGAGGCGCTTCGTCTACGGAGCGA
