## Supplementary material for "Stability of *orb* mRNA in *Drosophila* ovaries is regulated by *cis*-regulatory elements in the 3’ UTR and influences germ cell specification": File S2

**Amino acid Orb sequences**

Orb (915 AA)

MPLLQQYDTPDCSGSGGNMRALSGGSTTELLQKHSISSYLDHHHQQQQQQQHHLQLQQHQQQHSLLERCNDDGLISFINDPITLNDLLGLCGASTANEVVGTGQTPSTSAPILGAGGGGRANGVTAGAATATGVGVGAGGTLPGPGVPSIQGGGGGGVVGQQTNASCNTSAANPSASFGGNGSSSDVNNLLLASAAAAAAAAAGDGAQLSANAAAYAPLTPSSTHSSASPGTKSNFDYFQFENVAQSNPLKAFQRTNISFDCSAPLSPSTPTSIYNRSFHSSPLVSDSSNSSSGIGLSMDSINMFYNQQQQQQQPEQQGYTSLGNSMGSGLGLSLANASTRSNSPESQNSSNSTTEQNLLDMINLLSVNSNKIPHQQQQQQQQQQQQQQQQQQNQQQLQVQQQHQLQQQFVNLNRNYEQQISANLGSQQHGFEHNGVGVGASSSGNENCFSQYNLENITSVDMELAKLQNLQRINTLKLLHAQAQQMPLINQLLQSYAGNAIGSVGGSNLGNLMSAGGSSLMTEMAGNVGGIITTNDGHLDRVAKFYKSSAALCDATCTWSGHLPPRSHRMLNYSPKVFLGGIPWDISEQSLIQIFKPFGSIKVEWPGKEQQAAQPKGYVYIIFESDKQVKALLSACVLQVDDSHCGRNYFFKISSRRIKSKDVEVIPWIIADSNFVRSSSQKLDPTKTVFVGALHGKLTAEGLGNIMDDLFDGVLYAGIDTDKYKYPIGSGRVTFSNFRSYMKAVSAAFIEIRTTKFTKKVQVDPYLEDALCSICGVQHGPYYCRELSCFRYFCRSCWQWQHSCDIVKNHKPLTRNSKSQSLVGIGPSSSNVSLPFSGQRSIRDNRMGNGQHQQHQQHQYQQQKHRQLQEYSQPHSLNVMGNSGAANAAATSMVTLQQRQIHKVRIQRQQHQAI

Shortened Orb (840 AA)

MPLLQQYDTPDCSGSGGNMRALSGGSTTELLQKHSISSYLDHHHQQQQQQQHHLQLQQHQQQHSLLERCNDDGLISFINDPITLNDLLGLCGASTANEVVGTGQTPSTSAPILGAGGGGRANGVTAGAATATGVGVGAGGTLPGPGVPSIQGGGGGGVVGQQTNASCNTSAANPSASFGGNGSSSDVNNLLLASAAAAAAAAAGDGAQLSANAAAYAPLTPSSTHSSASPGTKSNFDYFQFENVAQSNPLKAFQRTNISFDCSAPLSPSTPTSIYNRSFHSSPLVSDSSNSSSGIGLSMDSINMFYNQQQQQQQPEQQGYTSLGNSMGSGLGLSLANASTRSNSPESQNSSNSTTEQNLLDMINLLSVNSNKIPHQQQQQQQQQQQQQQQQQQNQQQLQVQQQHQLQQQFVNLNRNYEQQISANLGSQQHGFEHNGVGVGASSSGNENCFSQYNLENITSVDMELAKLQNLQRINTLKLLHAQAQQMPLINQLLQSYAGNAIGSVGGSNLGNLMSAGGSSLMTEMAGNVGGIITTNDGHLDRVAKFYKSSAALCDATCTWSGHLPPRSHRMLNYSPKVFLGGIPWDISEQSLIQIFKPFGSIKVEWPGKEQQAAQPKGYVYIIFESDKQVKALLSACVLQVDDSHCGRNYFFKISSRRIKSKDVEVIPWIIADSNFVRSSSQKLDPTKTVFVGALHGKLTAEGLGNIMDDLFDGVLYAGIDTDKYKYPIGSGRVTFSNFRSYMKAVSAAFIEIRTTKFTKKVQVDPYLEDALCSICGVQHGPYYCRELSCFRYFCRSCWQWQHSCDIVKNHKPLTRNSKSQSLVGIGPSSSNVSLPFSGQ

Amino acid sequences used for polyclonal antibody generation are marked Yellow.
